# Wearable, High-Density, Time-Domain Diffuse Optical Tomography Array for Functional Neuroimaging

**DOI:** 10.1101/2025.04.22.649003

**Authors:** Kevin Renehan, Jinwon Kim, Heyu Yin, Fay Wang, Petros D. Petridis, Jack Grinband, Stephen H. Kim, Andreas H. Hielscher, Kenneth L. Shepard

**Affiliations:** Department of Electrical Engineering, Columbia University, New York, NY 10027, USA; Department of Biomedical Engineering, Columbia University, New York, NY 10027, USA; Department of Neurological Surgery, Columbia University, New York, NY 10027 USA; Department of Psychiatry, NYU Langone Center for Psychedelic Medicine, NYU Grossman School of Medicine, New York, NY, USA.10016 USA; Departments of Psychiatry and Radiology, Columbia University Vagelos College of Physicians & Surgeons, New York, NY, 10032 USA; Department of Radiology, New York University - Grossman School of Medicine, New York, NY 10010 USA; Department of Biomedical Engineering, New York University - Tandon School of Engineering, New York, NY 11201 USA

## Abstract

Noninvasive functional neuroimaging techniques, such as functional magnetic resonance imaging (fMRI) and electroencephalography (EEG), are essential tools for understanding brain activity and cognition for various neurological and mental health conditions. While fMRI offers high spatial resolution, its limited temporal resolution and costly large-form-factor restricts its accessibility and practicality for many applications. In contrast, EEG is more affordable and portable but has limited spatial resolution. In the present study, we overcome the limitations of existing neuroimaging technologies with the development of Micro-DOT, a functional near-infrared spectroscopy (fNIRS) system capable of high-density, time-domain diffuse optical tomography (HD-TD-DOT). Micro-DOT tackles the tradeoff between form factor and spatial resolution that has been a longstanding issue with existing fNIRS systems through the use of a unique hardware architecture that arrays HD-TD-DOT-capable electronics directly at the tissue surface. This is made possible with complementary-metal-oxide-semiconductor (CMOS) source-detector chiplets that contain all the electronics and optics necessary for HD-TD-DOT operation, and can be mounted on flexible polyimide packaging with a very minimal footprint. Pairing these hardware innovations with an advanced volumetric reconstruction software, Micro-DOT achieves in-plane spatial resolution, depth resolution, and localization accuracy comparable to fMRI, while maintaining the wearable form factor and portability of EEG, making it a viable stand-alone system for measuring subject-specific brain activation.

## INTRODUCTION

Functional neuroimaging techniques such as functional magnetic resonance imaging (fMRI) and electroencephalography (EEG) have well-established limitations. fMRI offers excellent spatial resolution (1-3 mm) but has poor temporal resolution (3-5 seconds) and requires a large-form-factor and costly instrumentation needed for superconducting magnets^1^. Additionally, fMRI is contraindicated for patients with metallic implants (such as cochlear implants and pacemakers), claustrophobia, movement disorders, and other conditions. Even for eligible patients, lying motionless during image acquisition can be uncomfortable. In contrast, EEG is performed with low-cost, wearable instruments and offers excellent temporal resolution (<10 ms) but limited spatial resolution (>5 mm)^2,3^.

Functional near-infrared spectroscopy (fNIRS) is an alternative to EEG and fMRI that makes use of backscattered near-infrared and infrared light for the purpose of measuring changes in the optical properties of the underlying tissue associated with the blood-oxygen-level-dependent (BOLD) response^4^. Similar to EEG-based devices, the relatively low cost, low complexity, high temporal resolution, and wearable form factor has underscored the potential of fNIRS-based devices to become portable tools for interrogating functional brain activity^5^.

However, fNIRS systems with in-plane spatial resolution of better than 1 cm generally have large form factors and high cost^2,6,7^. Wearable and low-cost fNIRS systems, on the other hand, have sparse source-detector (SD) arrangements and are limited by poor spatial resolution (> 1 cm) and non-existent depth resolution, producing only topographic maps of superficial brain activity^5,8^. For wearable fNIRS devices to become useful for subject-specific^9^ brain imaging, in-plane spatial resolution must be improved, and through-plane (or depth) resolution must be added^10^ without compromising on form factor.

With the goal of achieving high in-plane spatial resolution and depth resolution in a wearable device, we created the Micro-DOT system (**Fig. 1A**). Micro-DOT directly addresses the in-plane spatial resolution and depth resolution limitations of other wearable fNIRS systems through HD-TD-DOT imaging, a combination of both time-domain (TD-DOT) and high-density (HD-DOT) diffuse optical tomography. Diffuse optical tomography (DOT) is an extension of fNIRS that can generate three-dimensional, volumetric maps of BOLD changes in underlying tissue^11^ . In TD-DOT, one measures both the photon intensity at each detector, as in continuous-wave DOT (CW-DOT), and the arrival time, or time-of-flight, of incoming photons, which has been shown to increase contrast^12^ as well as improve in-plane spatial resolution and depth resolution in image reconstructions^13,14,15^. HD-DOT drives in-plane spatial resolution improvements through the arrangement of SD pairs, usually referred to as channels, at high density, where each source has a nearest detector that is less than 15-mm away^2^. This leads to more comprehensive coverage of the field-of-view with many overlapping measurements^2,16,17^. Through the combination of TD-DOT and HD-DOT, it has been shown that it’s possible to realize in-plane spatial resolutions of 1 mm^3^, rivaling that of fMRI, but these results come from large benchtop systems as opposed to small wearables^18^. The primary challenge in building TD-DOT wearables has been their relative complexity compared to CW-DOT wearables, with TD-DOT wearables requiring more advanced electronics capable of generating sharp optical pulses and timestamping incoming photons, which can contribute significant bulk^17,19^. In a similar way, the challenge of building HD-DOT wearables has been the need for additional channels, which also drives up the complexity, cost, and bulk^17,20^. Because of these challenges, many fNIRS wearables implement either TD-DOT or HD-DOT. In contrast, Micro-DOT implements both TD-DOT and HD-DOT in a wearable form factor by leveraging compact and scalable time-domain channel hardware that facilitates high density integration without adding significant cost or bulk.

**Fig. 1.**
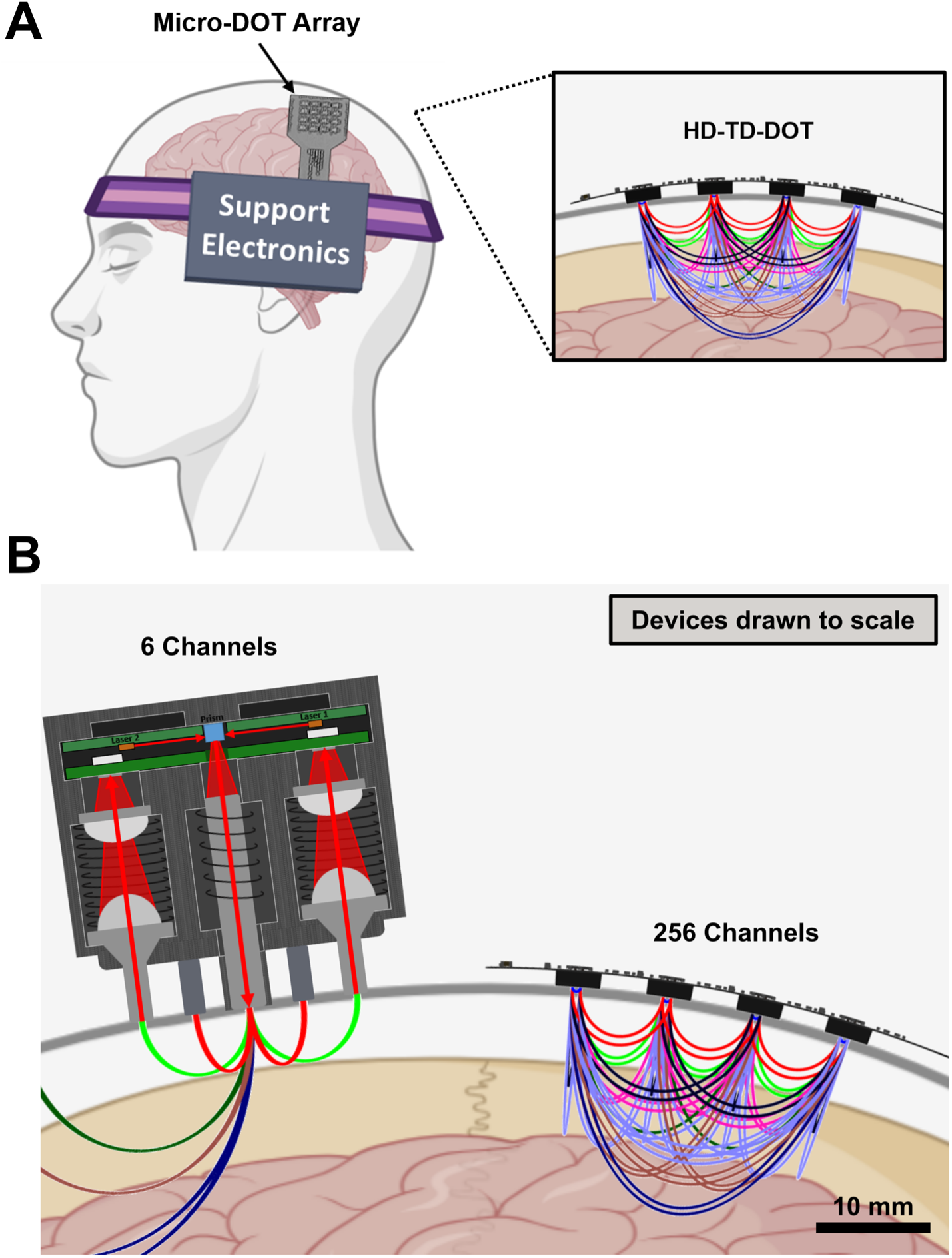
Comparison of TD fNIRS device form factors. **(A)** The form factor of the micro-DOT system showing the flexible array and supporting electronics mounted on the head. The micro-DOT array enabled HD-TD-DOT imaging, where time-domain channels are integrated at very high density to maximize spatial resolution in the FoV. **(B)** On the left, a commercial fNIRS module^23^ with light pipes incorporated for sources and detectors realizes six TD channels in the displayed volume. Heatsinks are included for the sources, as well as a prism for redirecting light towards the tissue. More channels are created if multiple modules are used, as is illustrated by the channels running off the left side of the image. The Micro-DOT device shown on the right lacks light pipes and uses less powerful sources that do not require heatsinking or additional optical components like prisms. Compared to the commercial module, which only includes six TD channels, Micro-DOT realizes 256 channels in a similar tissue area with approximately 10x less volume.

This paper begins with an introduction to the Micro-DOT’s unique hardware architecture, which is key to enabling the advancements that are presented here. What follows is a full validation of the Micro-DOT system, first at the source and detector level, and then at the system level. We evaluate the Micro-DOT system as a DOT imager using well-established protocols before extending the analysis to highlight improvements over low-density CW approaches in spatial resolution, localization, and imaging depth. We perform reconstructions in physiologically relevant optical phantoms to demonstrate the improvements in spatial resolution driven by the higher SD density that is supported by our form factor, while showing the spatial resolution improvements, especially at larger depths, provided by TD information. We conclude the work by presenting three-dimensional volumetric reconstructions in human subjects collected during three different experimental paradigms. After an initial occlusion experiment with the device positioned on the forearm, we conduct two brain-imaging studies: a simple breath-hold experiment with the device positioned on the forehead and a finger tapping experiment with the device positioned on the scalp above the primary motor cortex in which reconstructions can be compared with those resulting from fMRI.

## RESULTS

### Micro-DOT device architecture

The majority of existing fNIRS devices use light pipes or optical fibers as a way to couple sources and detectors to the tissue (**Fig. 1B**). This approach has several advantages: it provides thermal and mechanical isolation of sources, detectors, and associated electronics from the tissue; it can enable flexible reconfiguration of SD locations for optimization of channel density versus field of view (FoV); and more importantly it can achieve good coupling through hair with the light pipes acting as small wedges. This approach has several disadvantages as well: light pipes and fibers tend to be lossy, meaning that source power and detector light collection efficiency are sacrificed; the size of pipes and fibers, as well as other accessory optical components, largely limit the channel density; and the additional optical components add bulk and complexity.

Micro-DOT eliminates light pipes and optical fibers, and instead integrates high bandwidth sources and time-resolved detectors directly at the tissue interface. This results in better light collection efficiency at the detector and better preservation of source power while reducing weight and bulk in the system. Less powerful light sources can be used to achieve the same contrast-to-noise-ratio (CNR) performance, which is further improved with TD operation. We leverage these factors to greatly increase SD density (**Fig. 1B**) and improve image quality in Micro-DOT. A comparison of the Micro-DOT system to other state-of-the-art TD fNIRS systems^21,22,23^ is included in **Table S1**. **CMOS SD chiplet design.** The key building block for Micro-DOT is a 2.3 × 2.3 mm^2^ SD CMOS chiplet, an earlier version of which was described elsewhere^24^. Fabricated in a 130-nm high-voltage CMOS process, the chiplet includes an 8×8 array monolithic single-photon avalanche diode (SPAD) detectors for low-light, time-resolved detection of incident photons combined with two commercially available chiplet-mounted vertical cavity surface emitting lasers (VCSELs) at 680nm and 850nm. SPAD structures were chosen after a comprehensive design-of-experiments in which different parameters in the SPAD structure were varied (**Supplementary Section S1**). These laser wavelengths allow for detection of changes in the concentration of oxygenated hemoglobin (HbO2) and deoxygenated hemoglobin (HbR) in tissue^25^. The use of this CMOS chiplet architecture allows the SPADs, VCSELs, and all of the circuitry required for HD-TD-DOT to be fully integrated (**Fig. 2A,B**). Further details on ASIC implementation can be found in Methods, and a hardware block diagram is presented in **Fig. S1**.

**Fig. 2.**
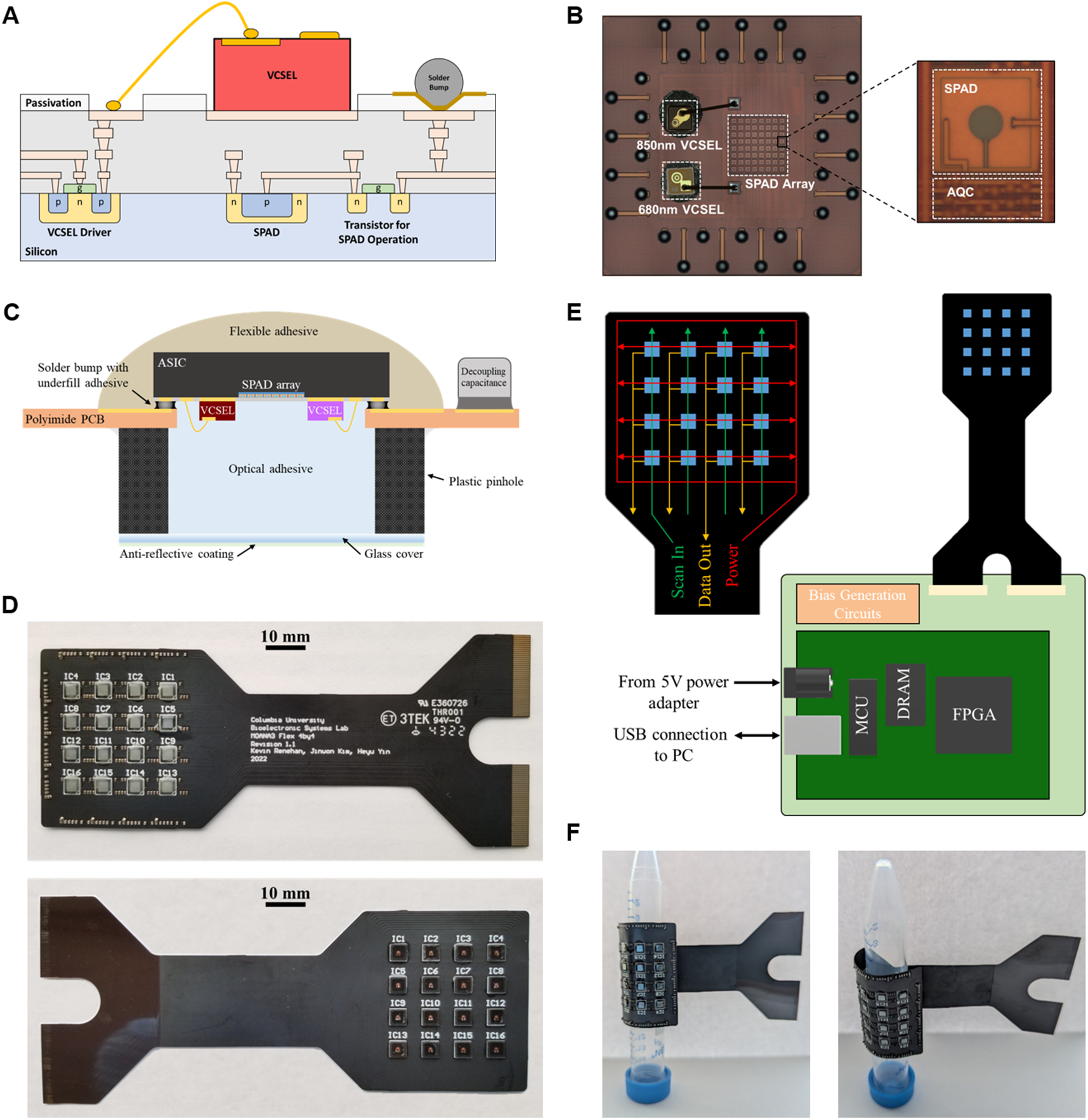
Flexible array packaging and system integration. **(A)** Cross-section of CMOS chiplet showing integration of monolithic SPAD detectors and top-side-bonded VCSEL sources directly alongside the analog and digital integrated circuits required for their operation. **(B)** Top-view of the 2.3×2.3 mm^2^ CMOS chiplet showing the relative positioning of the 680-nm VCSEL, 850-nm VCSEL, and SPAD array, as well as a close-up of a single SPAD and the active quench and reset circuits (AQC). **(C)** Cross-sectional schematic of CMOS chiplet flip-chip bonded to a flexible PCB with optical components positioned to couple to the tissue through a small cutout in the polyimide substrate. **(D)** HD-TD-DOT array with CMOS chiplets arrayed in a 4×4 grid layout on an 8-mm pitch. **(E)** Illustration of signal organization on the flexible array facilitated by tileable chiplet design and the arrays’ connection to supporting electronics that handle bias generation, data collection, and communication with a host PC. **(F)** Micro-DOT array wrapped around a plastic centrifuge tube with an 8-mm radius and secured with a piece of double-sided tape to demonstrate the flexibility of the device.

### Array design

Each chiplet is capable of functioning as a standalone TD fNIRS imager. In Micro-DOT, they are tiled into an array for HD-TD-DOT imaging on a flexible printed circuit board (PCB) as shown in **Fig. 2D**. This board contains 16 SD chiplet locations in a 4×4 grid with a center-to-center distance between adjacent SD locations of 8 mm^25^. With each chiplet hosting a single SPAD array and two VCSELs, the total number of sources and detectors on the array is 32 and 16, respectively. To facilitate tiling within the array, configuration signals are shared row-wise and power rails are shared column-wise, with each chiplet requiring only two unique signals (**Fig. 2E**). A shared clock is routed across the array in an H-tree structure such that all chiplets are synchronized to the same global timing reference. These interconnections require only two metal layers in the PCB, allowing its thickness to remain below 110 µm. The array consumes 965 mW in total during normal imaging operation.

Chiplets are solder-bump attached to the PCB with a laser cutout in the PCB exposing the SPAD array and VCSELs, allowing the source and detector arrays to be in close proximity to the tissue for maximal light collection efficiency (see Methods). A square pinhole (**Fig. S2**) with a 4-mm side length is placed over each cutout to eliminate light leakage paths from chiplet to chiplet, while also protecting the surface of the chiplet and the VCSELs from damage and thermally isolating the electronics on the array from the tissue. These pinholes also provide some ability to wedge through short hair (**Fig. S3**). The VCSELs operate multi-mode with a peak emission angle of approximately 10 degrees, which is unaffected by the pinhole. On the topside of the array, which faces away from the tissue interface, a thermally conductive, flexible epoxy protects the backside of the chiplet and also enlarges the backside surface area for heatsinking. The fully assembled array can support a bending radius of better than 8 mm (**Fig. 2F**), which allows for imaging of structures with significant curvature such as the fingers. A cross-section of the array stackup is shown in (**Fig. 2C**). The assembly steps (see Methods) required to fabricate the Micro-DOT array are presented in **Fig. S4**.

### System Integration

The array connects directly to a motherboard (**Fig. S5**) that hosts the peripheral electronics needed for proper operation (see Methods), including bias generation circuits and an Opal Kelly XEM7010 module containing a field-programmable gate array (FPGA), dynamic random-access memory (DRAM), and a USB-capable microcontroller (MCU) (**Fig. 2E**). Data from all 16 ASICs is aggregated on the FPGA (**Fig. S6**). The system communicates via USB 2.0 with a host PC, where data can be visualized in real-time using a custom Python GUI (**Fig. S7**) and simultaneously stored for later analysis. The fully assembled flexible array weighs only 2.5 grams, while the entire Micro-DOT system, including the flexible array and motherboard, weighs 90 grams in total. The system consumes 3.4 W power in total during imaging.

### Single SD channel validation

Prior to using the system for full HD-TD-DOT reconstructions, we characterize single SD channels using the basic instrumental performance (BIP) protocol^26^ and the nEUROPt protocol^27^. BIP characterizes performance metrics including detector responsivity, measurement stability, and the impulse response function (IRF). A complete summary of BIP metrics is presented in **Fig. S8**. In particular, the IRF of the system, dominated in the early portion by the width of the VCSEL pulse and in the later portion by the SPAD’s diffusion tail, is a critical metric for determining how accurately a photon’s time-of-flight can be measured and has significant impacts on contrast for deeper inclusions. The IRF for Micro-DOT, as measured in reflectance mode (**Fig. 3A**), is shown in **Fig. 3B**, demonstrating a full-width at half-maximum (FWHM) of 505 ps and 706 ps for the 680-nm and 850-nm wavelengths, respectively. The more important full-width at 1/1000-maximum (FW1/1000M) is given by 7.36 ns and 8.8 ns for 680 nm and 850 nm (**Fig. S9**), respectively, reflecting the diffusive tails in the response of the SPADs as determined by the depth of the multiplication region and the thickness of the neutral regions within the SPAD structure.

**Fig. 3.**
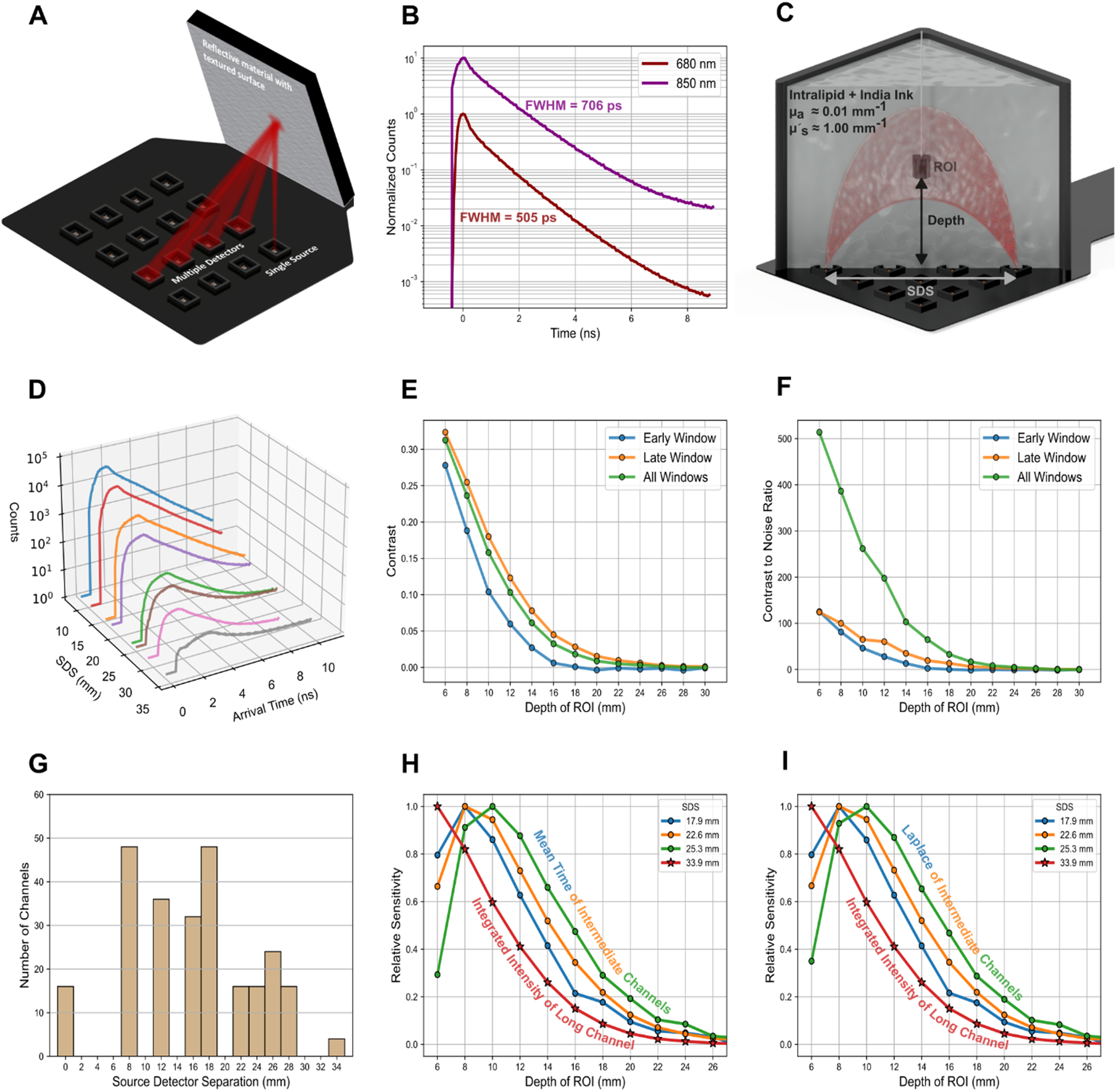
Electrical and optical array characterization. **(A)** Test setup for measuring the array IRF from a single source location to various detector locations on the array. **(B)** Example IRF of 680 nm source (normalized to 1) and 850 nm source (normalized to 10) for a channel with 8-mm SDS. **(C)** Test setup for collection of optical characteristics as part of the nEUROPt protocol with black plastic cylinder positioned at variable depths above the array and submerged in a scattering media. **(D)** ToF histograms as a function of source-detector separation for 28.89-mm channel in the nEUROPt setup. **(E)** Contrast results for the 28.89-mm channel. **(F)** Contrast-to-noise results for 28.89-mm channel. **(G)** Histogram of channel separations for the array. **(H)** Comparison of the sensitivity of the integrated intensity parameter from a 33.9-mm channel and the mean time parameter from three channels with shorter SDS values. **(I)** Comparison of the sensitivity of the integrated intensity parameter from a 33.9-mm channel and the Laplace parameter from three channels with shorter SDS values.

The nEUROPt protocol^27^ characterizes metrics such as contrast, contrast-to-noise, spatial resolution, and depth selectivity. To measure these parameters, a custom setup (see Methods) was built using a 5 × 5 × 3 cm^3^ tank, filled with a scattering and absorbing liquid with a reduced scattering coefficient of µs’ = 1.0 mm^-1^ and absorption coefficient of µa = 0.01 mm^-1^, mounted directly onto the surface of the array (**Fig. 3C**). A dark-black, 100 mm^3^ cylinder (5 mm diameter, 5 mm height) forms a region-of-interest (ROI) suspended at a particular distance from the SD plane at the lateral center point of the array. The ROI can be moved along the z-axis from a depth of 6 mm to 30 mm. Time-of-flight (ToF) histograms (**Fig. 3D**) are collected for selected channels with and without the presence of the ROI for an integration time of one second. Contrast (**Fig. 3E**) and CNR (**Fig. 3F**) results at 680 nm are shown for a single channel with 28.84 mm SD separation (SDS) in an early window (0.0 to 0.5 ns), late window (2.0 to 2.5 ns), and for all windows (0.0 to 10.0 ns) of the ToF histogram. As expected, the late window gives the highest contrast for deeper inclusions. However, the late window also presents higher contrast for shallower inclusions because of the system’s relatively broad IRF, which redistributes early photons into later time bins. Contrast and CNR results for other SDSs are presented in **Fig. S10**.

### Feature extraction from the ToF histograms

Rather than using explicit time windows as is done in the nEUROPt protocol, we instead analyze ToF histograms by extracting temporal features through the Mellin and Laplace transforms. From the Mellin transform, we can determine the statistical moments – here, we focus on the mean time (*M*). From the Laplace transform (Laplace feature, *L*), we can “gate” the ToF histogram (see Methods) to isolate characteristics of the late photons arriving from deeper tissues^28,29,30^. When employed in reconstruction algorithms, these features have been shown to improve overall image quality and convergence time^18,31^. These features have also been shown to be more resilient to imperfect IRFs compared to explicit time windows while still providing higher sensitivity to contrast at depth than CW information alone^32,33^. In addition to these two temporal features, we also extract integrated intensity (*E*) as would be done for a CW device.

### HD-TD-DOT capabilities

When all 16 source locations and 16 detector locations are considered, a total of 256 channels are measured in each imaging frame with a variety of SDSs (**Fig. 3G**). In CW devices, it has been suggested that the optimum SDS for probing the brain is between 30 mm and 35 mm, driven by enhanced sensitivity for the cortical tissue as SDS is increased^34^. Channels with short SDS, less than 10 mm, are optimal for measurement of responses in shallow tissues, such as the scalp, that can be regressed out of long-channel measurements^35^. The utility of channels with SDSs between 10 mm and 30 mm is not as well defined for CW brain imaging devices. In contrast, our HD-TD-DOT array has a large proportion of channels (73.4%) with SDS between 10 mm and 30 mm. An important distinction between a TD device and a CW device is the ability for channels having intermediate (and even short) SDS to still be useful in the interrogation of deeper tissue^32,36^.

The importance of shorter SDS channels in providing useful depth information can be demonstrated within the framework of the nEUROPt protocol by extending the analysis to include changes in *M* and *L* features in response to the presence of the ROI at different depths. In particular, we measure relative sensitivity^33^, in which the distribution of changes in the *E*, *M*, and *L* features as a function of depth are normalized to the corresponding peak values to assess how strongly these features respond to superficial activation compared with deeper activation. We find that the relative sensitivity of the *M* and *L* features extracted from a channel with a SDS as short as 17.89 mm exceeds the relative sensitivity of the *E* feature extracted from a channel with an optimal SDS of 33.9 mm (**Fig. 3H,I**). This is critical to the operation of the Micro-DOT device; the sensitivity of shorter channels to changes in deeper layers means that increasing the number of channels in a given area can lead to improvements in spatial resolution. Our TD array has 124 channels with SDS of 17.89 mm or longer that are all capable of detecting activation at depths greater than 15 mm; in contrast, a CW array of the same extent would have only 20 channels capable of measuring the same depth (channels with SDS > 28 mm). TD operation is, therefore, key to realizing HD brain imaging in this form factor. nEUROPt results for other SDSs, as well as mean-time-to-noise ratio (MNR), Laplace-to-noise ratio (LNR), and lateral spatial resolution results are included in **Figs. S10-S12**.

### *In vitro* reconstructions of brain phantom

To validate the Micro-DOT system and the associated image reconstruction algorithms with known ground truth, we created a phantom to mimic the optical properties of the cortex and the layers of tissue above it. The optical properties, as well as the thicknesses of the scalp, skull, cerebrospinal fluid (CSF), and grey matter (GM) layers in the phantom, were chosen to be consistent with tissue beneath the Oz landmark in the 10-10 system for EEG^35^. Optical properties of each layer are presented in **Table S2**. A dark black ROI of the same material and with the same cylindrical shape and optical properties as the one used in the nEUROPt protocol was suspended at various positions and depths in the GM layer to mimic an absorptive change due to neural activation (**Fig. 4A**). The percentage changes in E, M, and L features as extracted from the ToF histograms at each detector location are presented for different source and ROI positions, as shown in **Fig. 4B**. The ability of the imager to localize a 100 mm^3^ cylindrical ROI (5 mm diameter, 5 mm height) within the GM layer was evaluated by calculating the localization error and spatial resolution following image reconstructions using E, M, and L features for 1 s of integration time per source. Image reconstructions were performed using the SENSOR code,^18^ which provides for ultrafast and ultrahigh-resolution DOT. With the ROI positioned at a depth of 16 mm, E, M, and L features localize the ROI with only ∼1.5mm error (**Fig. 4C**, top row); at 20 mm, the improvement in localization using M and L becomes apparent when compared to the use of E alone (**Fig. 4C**, middle row). Two 21-mm^3^ ROIs (3 mm diameter, 3 mm height) separated edge-to-edge by 3 mm at a depth of 16 mm (**Fig 4C**, bottom row) are resolved in the M and L reconstructions but are not in the E reconstruction, clearly demonstrating that TD information can improve spatial resolution compared to CW techniques when imaging deep cortical tissue.

**Fig. 4.**
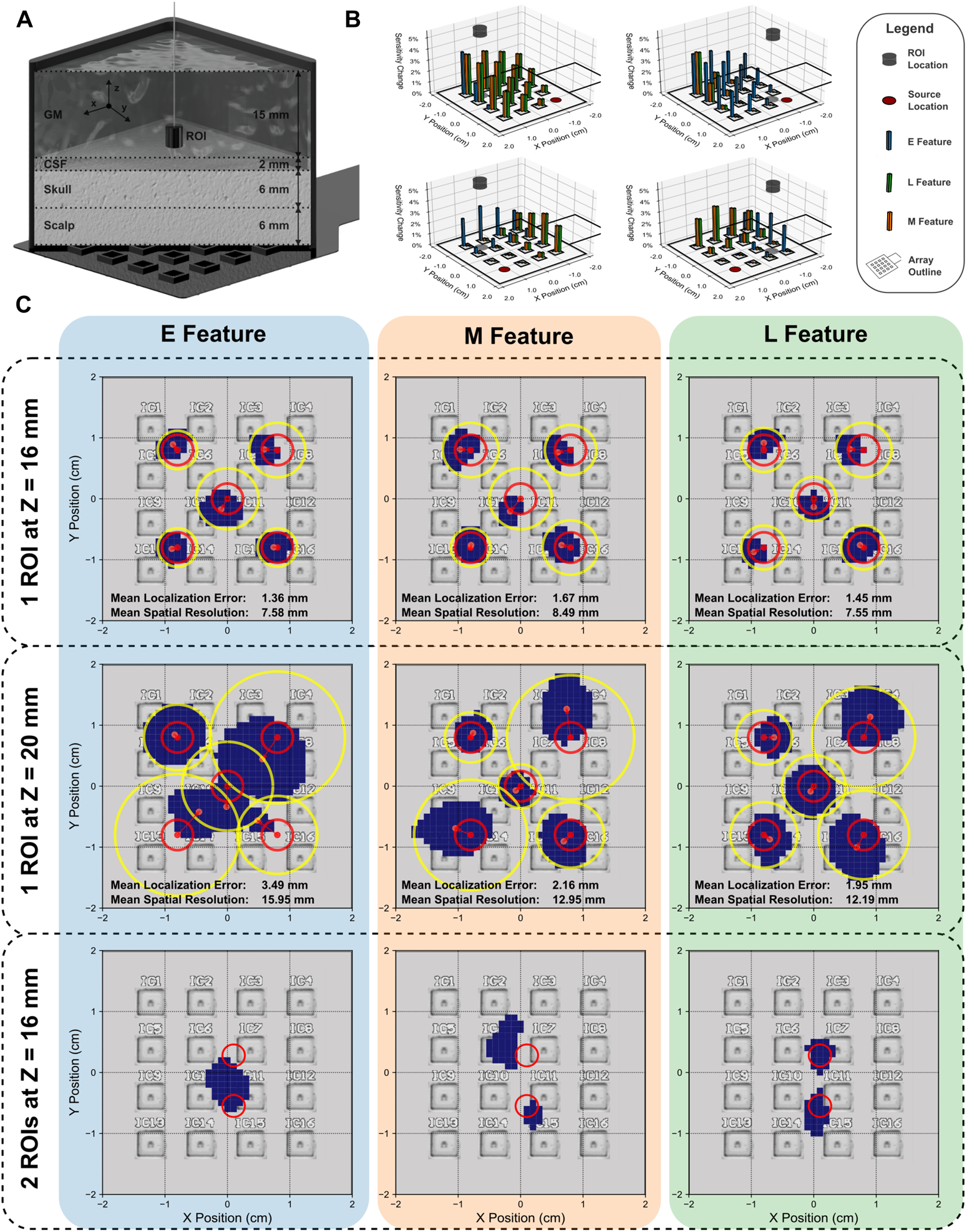
Image reconstruction in a brain phantom. **(A)** Setup for brain phantom image reconstruction experiments showing approximate thicknesses of each layer. **(B)** Visualization of the changes in *E*, *M*, and *L* features as source position on the array and ROI position within the GM layer are varied. The ROI is shown above the array and is also projected to the Z = 0 plane to indicate its XY position. **(C)** Image reconstruction results using *E*, *M*, and *L* features for five single ROI experiments at 16-mm depth (top row), five single ROI experiments at 20-mm depth (middle row), and a dual ROI experiment at 16-mm depth (bottom row). Red circles indicate ground truth ROI locations. Localization error is defined as the length of the red arrows pointing from the centroid of the reconstructed target to the center of the ground truth. Spatial resolution is defined as the diameter of the yellow circle, which is concentric to the red circle defining the ground truth ROI location and completely encloses the reconstructed target.

### *In vivo* imaging of the human forearm and forehead

For the first *in vivo* demonstration, we conducted a venous occlusion experiment on the forearm and a breath-hold experiment on the forehead. Venous occlusion was performed with a blood pressure cuff (BPC) placed just above the elbow. The protocol consisted of a two-minute baseline, followed by one minute of occlusion, and concluded with three minutes of recovery (**Fig. 5A**). The vasculature on the underside of the forearm is immediately visible allowing us to position the device over a prominent vein (**Fig. 5A, inset**). During the occlusion period, the BPC was inflated to a pressure of 60 mmHg.

**Fig. 5.**
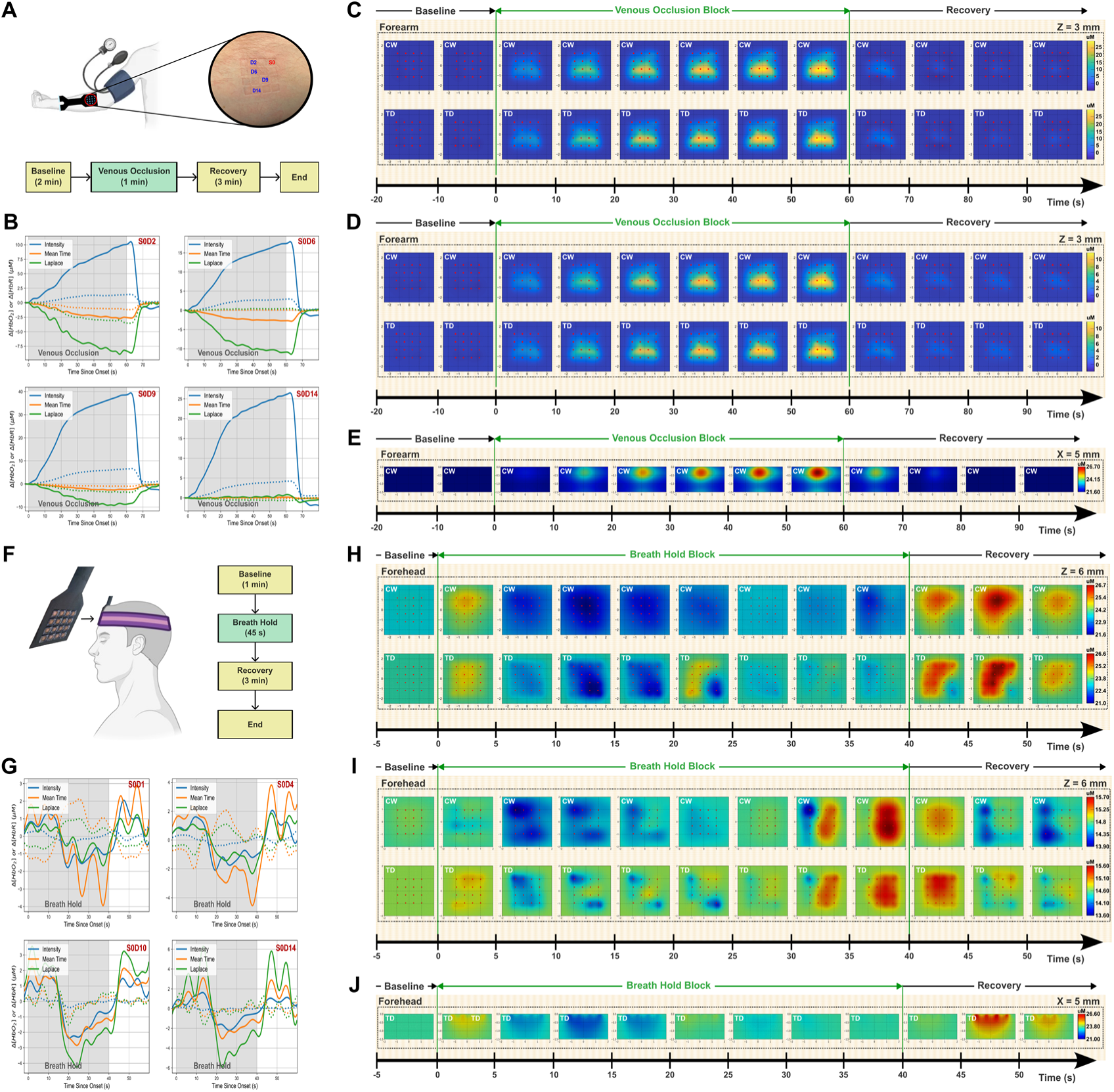
Forearm and forehead imaging results. **(A)** Illustration of the experimental setup and schematic diagram of the experimental protocol for the forearm venous occlusion experiment. Array imprints on the tissue surface show positioning of device relative to vasculature. **(B)** Block averages of HbO2 (solid lines) and HbR (dashed lines) changes resulting from E, M, and L features in response to the 1-min venous occlusion task at various SDS. **(C)** Image reconstruction results showing changes in HbO2 concentration for an XY plane at Z = 3 mm with CW and TD results plotted for comparison. **(D)** Image reconstruction results showing changes in HbR concentration for the same XY plane at Z = 3 mm. **(E)** Image reconstruction results changes in HbO2 for a YZ cross section at X = 5 mm. **(F)** Illustration of the experimental setup and schematic diagram of the experimental protocol for the forehead breath hold experiment **(G)** Block averages of HbO2 (solid lines) and HbR (dashed lines) changes resulting from E, M, and L features in response to the 1- min venous occlusion task at various SDS. **(H)** Image reconstruction results showing changes in HbO2 concentration for an XY plane at Z = 6 mm with CW and TD results plotted for comparison. **(I)** Image reconstruction results showing changes in HbR concentration for the same XY plane at Z = 6 mm. **(J)** Image reconstruction results showing changes in HbO2 for a YZ cross section at X = 5 mm.

Block averages of *E*, *M*, and *L* revealed large changes that were temporally consistent with the onset and offset of occlusion (**Fig. 5B**). The occlusion has the most significant impact on *E*, which is very sensitive to shallow oxygenation changes. Image reconstructions were carried out with the multispectral PDE-constrained inverse model. Both CW- and TD-DOT image reconstructions of HbO2 (**Fig. 5C**) and HbR (**Fig. 5D**) for an x-y plane at a depth of 3 mm show a strip of activity consistent with the location of the vein through the occlusion period. CW-DOT reconstructions of HbO2 for a y-z plane at x = 5 mm show the cross-section of the vein, with the center of the vein located at a depth of approximately 3 mm (**Fig. 5E**). The temporal behavior of the spatial reconstructions closely aligns with that of the block averages.

The breath-hold experiment was designed with the subject sitting upright to maintain natural posture and minimize motion artifacts with the Micro-DOT device positioned in the center of the forehead between the top of the eyebrows and the bottom of the hairline. Following a 1-minute baseline, the breath-hold was conducted as a 40-second block followed by three minutes of recovery (**Fig. 5F**). CW- and TD-DOT image reconstructions of HbO2 (**Fig. 5H**) and HbR (**Fig. 5I**) for an x-y plane at a depth of 6 mm show consistent features that align with expected physiology. Over the course of the breath-hold, HbO2 levels decrease; HbR levels, after an initial decrease, reach a peak by the end of the breath-hold. During the recovery phase, an overshoot of HbO2 and HbR relative to the baseline is observed. These responses are consistent across different SDS for both CW and TD (**Fig. 5G**). TD-DOT reconstructions of HbO2 activity for a y-z plane at x = 5 mm show uniform changes in the first 6 mm of the tissue, consistent with the approximate thickness of the scalp, and fall off sharply at further depths that are likely associated with the skull (**Fig. 5J**).

### *In vivo* imaging of the motor cortex of the human brain

We performed block-design, right-hand-finger tapping experiments with the array positioned over the left motor cortex of a human subject. Finger tapping was conducted in 6-s blocks with 20 s of rest between sequential blocks for a total of 30 blocks (**Fig. 6A**). The subject was a bald, Caucasian male with Fitzpatrick skin type III. Prior to the experiment, fMRI was used to assess the location of the brain region that showed maximal motor cortex activity in response to the finger tapping task. The structural T1 and the activation maps were loaded into BrainSight to aid in positioning the Micro-DOT array on the scalp relative to the ROI (**Fig. 6B**). After array placement, the rest of the system was secured on the head with a Velcro strap and covered with a black shower cap to block unwanted ambient light. Scalp coupling index (SCI) was assessed after array placement to validate coupling of sources and detectors to the tissue^37^ (**Fig. 6C**). Block averages of changes in HbO2 and HbR through the duration of the task show a clear response that is consistent across the array for many different SDSs (**Fig. 6D**).

**Fig. 6.**
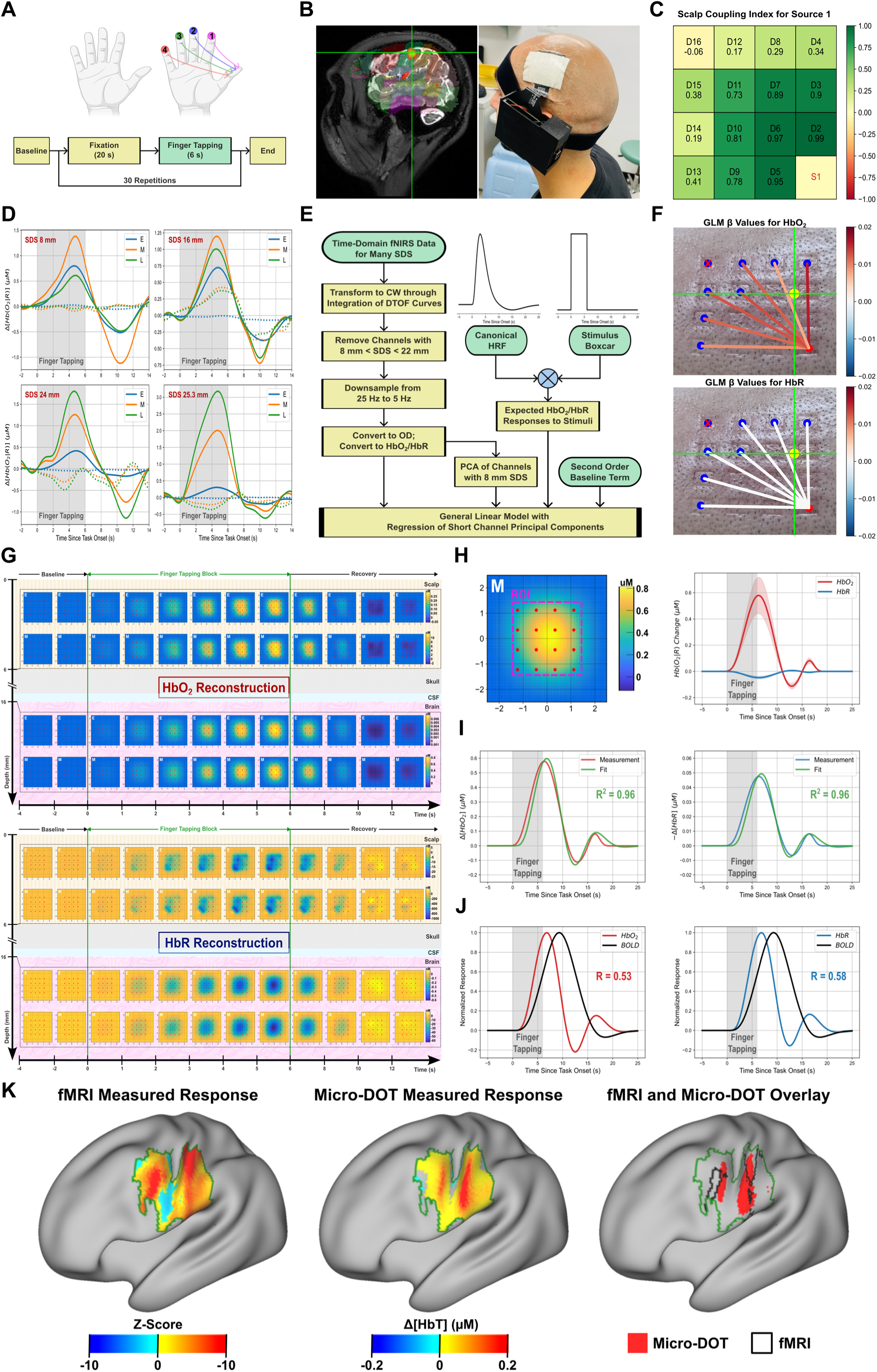
Motor cortex imaging results. **(A)** Finger tapping protocol. **(B)** fMRI localized activation during finger tapping task and corresponding array placement. Right panel depicts author Petros D. Petridis wearing the Micro-DOT system. **(C)** Scalp coupling index for a single source. **(D)** Block averages for four different channels of varied source-detector separation in response to task. **(E)** GLM data processing block diagram. **(F)** GLM results for HbO2 (upper) and HbR (lower). **(G)** Image reconstruction results showing changes in HbO2 (upper panel) and HbR (lower panel) concentrations 6 mm (scalp) and 16 mm (brain) below the array with CW and TD results plotted for comparison. **(H)** Normalized HbO2 and HbR temporal responses (right) in selected ROI shown on the left. **(I)** FLOBS fitting results for HbO2 (left) and HbR (right) with R^2^ values shown. **(J)** HbO2 (left plot, red trace) and HbR (right plot, blue trace) normalized HR derived from FLOBS coefficients compared to fMRI normalized HR with correlation coefficient shown. **(K)** Comparison of fMRI and Micro-DOT activation and deactivation in the left hemisphere in response to finger tapping task with the right hand. The comparison is cropped to show only the field-of-view of the Micro-DOT device, which is represented by the dark green outline in each of the images. A map of activation and deactivation as measured by fMRI is shown on the left. A map of ΔHbT as measured by Micro-DOT is shown in the middle. A comparison of strongly activated regions between fMRI and Micro-DOT is shown on the right. Strongly activated voxels are those falling in the 75^th^ percentile or above within their respective dataset. Strongly activated regions measured with Micro-DOT are shown in red, and the black outlined regions correspond to strongly activated regions measured with fMRI for comparison.

To show the relevance of our results to CW fNIRS systems, we transformed our HD-TD-DOT data into HD-CW-DOT data by considering only the *E* feature, and processed the resulting dataset in MATLAB using NIRS Brain AnalyzIR Toolbox^38^ with a GLM analysis pipeline^39^ (**Fig. 6E**). The response amplitudes, or β values, retrieved by the GLM (see **Supplementary Section S2**) were relatively consistent across the field-of-view, with β having a mean of 0.1171 µM and a standard deviation of 0.0254 µM (**Fig. 6F**).

Recent work has shown that GLM can be extended to consider TD moments, but the same work also suggests that image reconstruction incorporating TD analysis is a better mechanism for assessing TD-DOT data in the case of many overlapping channels^32^. Reconstructions were performed utilizing the multispectral SENSOR algorithm^18^, employing either *E*, *M*, or *L* as the selected feature. HD-CW- and HD-TD-DOT results for both HbO2 and HbR at 6 mm depth (scalp layer) and 16 mm depth (brain layer) are shown beginning 2 s before the finger tapping block and ending 4 s after the finger tapping block concludes (**Fig. 6G**). As expected, scalp activation at 6 mm depth is task evoked^40^.

To validate our results, we performed spatial and temporal comparisons between Micro-DOT data and measurements of the BOLD signal taken with fMRI (fMRI-BOLD). We extracted the hemodynamic responses for both HbO2 and HbR from a group of voxels over the duration of the task (**Fig. 6H**) and performed a fitting with three basis functions taken from FMRIB’s Linear Optimal Basis Sets (FLOBS). R-squared coefficients for the FLOBS fits were 0.96 and 0.96 for HbO2 and HbR, respectively (**Fig. 6I**). We then created a representative fMRI-BOLD hemodynamic response (HR) to the finger tapping task by convolving the task regressor with the individual subject’s fMRI-BOLD hemodynamic impulse response function (HRF) for comparison with the HR measured by Micro-DOT. The correlation coefficient (R) between the fMRI-BOLD HR and the Micro-DOT HR was 0.53 for HbO2 and 0.58 for Hb. Comparisons of the normalized responses of HbO2 and HbR to the fMRI-BOLD signal (**Fig. 6J**) reveal some interesting details. First, the temporal profiles of HbO2 and HbR are almost identical (when the inversion of HbR and the amplitude differences are not considered), despite the physiological expectation that HbR should peak after HbO2. Second, the correlation between HbR and fMRI-BOLD is only slightly higher than between HbO2 and fMRI-BOLD, despite the expectation that HbR should have much higher correlation to the fMRI-BOLD signal. Lastly, the temporal profile of HbO2 peaks about 2.4 s earlier than the fMRI-BOLD signal. All of these observations are consistent with those commonly reported in fNIRS literature^41^.

For spatial comparisons, we perform registration of the Micro-DOT ΔHbT reconstruction volume to the T1-weighted scan, transform the volumes into MNI space, and then perform volume-to-surface mapping in the left hemisphere such that direct comparisons can be made between fMRI and Micro-DOT. Unthresholded fMRI-BOLD and Micro-DOT activity are shown within the FoV of the Micro-DOT device (**Fig. 6K**, left and middle column) and show good agreement, with a Dice coefficient of 0.72 and both imaging methods reporting widespread activation in the FoV in response to the finger tapping task. To better localize the response to the finger tapping task, regions of strong activation were identified in each dataset by considering only those voxels with response amplitudes in the 75^th^ percentile or higher (**Fig. 6K**, right column). Both fMRI and Micro-DOT resolve two distinct regions of strong activation in similar locations, demonstrating that the Micro-DOT device is capable of properly localizing changes in brain oxygenation resulting from finger tapping to the precentral and postcentral gyrus. Compared to fNIRS systems with lower spatial resolution, Micro-DOT allows for more detailed localization of the activated brain regions.

## CONCLUSION

In this work, we engineered and validated Micro-DOT, an HD-TD-DOT array paired with a capable reconstruction backend targeted at pushing the current boundaries of spatial resolution, depth resolution, and localization accuracy in fNIRS in a wearable device. Our aggressive form factor is enabled by heterogeneous integration of all necessary optical and electronic components on an CMOS chiplet with a 2.3 x 2.3 mm^2^ footprint that can be tiled to form larger HD-TD-DOT arrays, while improvements in spatial resolution are realized by leveraging time domain moments^33^, TD-DOT^13^, HD-DOT^2^, and state-of-the-art image reconstruction algorithms^18^. We have shown the capability of the system to resolve 100 mm^3^ inclusions in physiologically relevant optical phantoms with localization error on the scale of a few millimeters at depths as large as 20 mm as well as to correctly distinguish multiple 21 mm^3^ inclusions separated by only 3 mm at a depth of 16 mm. In addition, we demonstrate that the system is capable of localizing brain activation mapped to underlying anatomical structures, providing new capabilities for fNIRS for clinical applications in which single-subject specificity is desired. These include many applications for which fMRI is the main diagnostic imaging tool, including assessing the extent of damage after stroke or traumatic brain injury; localization of seizure foci in epilepsy; diagnosis and monitoring of psychiatric disorders such as depression, anxiety, and post-traumatic stress disorder, particularly in response to treatment; and monitoring the progression of neurodegenerative diseases. For glioblastoma, this device could allow for outpatient monitoring of cerebral blood flow such that vascular dysregulation at the site of a resected tumor could be identified and used as a marker for tumor recurrence^42^. If the device is paired with a reconstruction backend that employs a neural network for improved convergence speeds^43^, image reconstruction could be performed in real-time. This could enable an entire class of experiments related to biofeedback that would have very minimal complexity compared to fMRI biofeedback setups^44^.

Limitations of the current design include challenges for optical coupling caused by the density and color of hair, though this is a problem for all optical imaging methods The current support electronics connected to the patch are still too bulky for an unobtrusive wearable; this can be considerably improved as well as made wireless and battery-powered. Although more than one device can be positioned on the head for applications in which simultaneous measurement of multiple brain regions is required, it may also be desirable to also expand the field-of-view (FoV) of each array. The advantage of the Micro-DOT chiplet architecture is that is easily scalable to different array sizes. However, if the same SD density is to be maintained in a larger FoV, it will result in a reduction in available integration time for each source, which will make maintaining signal-to-noise ratio (SNR) in the time domain moments even more challenging. These limitations can be largely addressed with simple enhancements to the existing design. Source power can be increased through the use of VCSEL arrays, which continue to improve in efficiency and density. Enhancements in photosensitive area are possible through realization of the chiplet in a more aggressive CMOS technology node. Both of these changes would increase the number of detected photons, which would lead to better SNR for a given integration time. Major improvements in the IRF of the SPADs, particularly in the FW1/1000M, are also possible in different technology nodes, which would lead to even better sensitivity in the time-domain parameters. It should also be mentioned that our form factor is very well-suited to integration of sensors for concurrent measurement of physiological signals such as motion, temperature, skin conductance, and oxygen saturation.

In summary, the system presented in this work represents a new type of diffuse optical tomography device that simultaneously addresses limitations in spatial resolution and wearability that have been a primary factor in preventing fNIRS from gaining acceptance as a stand-alone technique for measuring subject-specific brain activation.

## SUPPORTING INFORMATION

Supporting information includes Sections S1 and S2, Figures S1 through S32, Tables S1 through S3, and extended methods and analysis.

## DATA AND MATERIALS AVAILABILITY

All imaging data is available at https://github.com/klshepard/microDOT. All other relevant data are available from the corresponding authors upon reasonable request.

## CODE AVAILABILITY

All and all scripts used for data analysis and image processing are available at https://github.com/klshepard/microDOT. All other relevant data are available from the corresponding authors upon reasonable request.

## ACKNOWLEDGEMENT

This work was supported in part by Defense Advanced Research Projects Agency (DARPA) under Contracts N6600119C4020 and HR00112320037 (K.L.S) and by the National Science Foundation under Grant 2141006 (K.L.S). Fay Wang was supported by a National Science Foundation Graduate Research Fellowship under Grant No. DGE-2036197. We gratefully acknowledge TSMC for chip fabrication and their support in the use of experimental SPAD devices.

## AUTHOR CONTRIBUTIONS

K. R. and K.L.S. conceptualized the study. K.R., J.K., H.Y., and K.L.S. conceptualized the electronic and optical packaging, array design, and system design. K.R. designed the circuits. K.R. and J.K. designed the system. K.R. and J.K performed the BIP and nEUROPt tests and corresponding data analysis. K.R and J.K performed brain phantom experiments. K.R., J.K., and H.Y. performed the *in vivo* experiments. K.R. and J.K. performed GLM analysis for *in vivo* experiments. F.W., S.K., and A.H. performed image reconstruction for brain phantom experiments and all *in vivo* experiments and provided guidance for experimental design. J.G. and P.P. performed fMRI experiments and provided supervision and guidance for Micro-DOT brain imaging experiments. J.G. conceptualized fMRI and Micro-DOT spatial and temporal overlaps.

K.L.S. provided overall supervision and guidance.

## CORRESPONDING AUTHORS

Correspondence and requests for materials should be addressed to K.L.S.

## MATERIALS AND METHODS

### Design and fabrication of the Micro-DOT system

#### Design of the CMOS chiplet

The Micro-DOT chiplet has a 2.3 × 2.3 mm^2^ footprint and is fabricated in a TSMC 130-nm high-voltage CMOS process. Analog circuit design is done in Cadence Virtuoso and simulated with Cadence Spectre. Digital design is done with Synopsys Design Compiler (logic synthesis) and Cadence Innovus (floorplanning, place and route). Digital verification is done using Cadence Xcelium with results visualized in Cadence NC-Verilog. Top level verification is done with Calibre nmDRC (physical verification) and Calibre nmLVS (layout versus schematic verification). Rather than having source electronics that are separate from detector electronics, as is typical in commercial fNIRS devices, the chiplet combines both, creating both a source location and a detector location at each point in the array, driving up attainable SD density.

#### SPAD characterization

The chiplet includes an 8×8 array of SPADs. The technology node used for the chiplet includes a custom SPAD implant that has been optimized for sensitivity at NIR and IR wavelengths, with a cross section shown in **Fig. S2A**. When a single SPAD is tested individually, separate from the circuits on the chiplet, the SPADs have photon detection probability (PDP) of 18.05% and 11.5% at 670-nm and 850-nm, respectively. Dark count rate (DCR) for individual SPADs is 28.3 counts per second, and individual SPAD structures have a fill factor of 18.5%. This sets the SPAD’s noise equivalent power (NEP) to 0.06 fW/√Hz for the 680-nm wavelength, and 0.08 fW/√Hz for the 850-nm wavelength. The low NEP of SPADs is beneficial in DOT, since photon counts are low and readout noise must be minimized.

When SPADs are integrated into an array that includes active circuits, the fill factor reduces to approximately 4.84% based on the pixel size. The DCR also increases to 70 kHz due to the heat generated by active circuits. While we cannot directly measure all of the parameters required for an NEP calculation once the SPADs have been integrated into the array on the chiplet, we do have the ability to measure certain parameters and make assumptions about others. The PDP of the SPAD is assumed to be the same after integration into the array. The full SPAD array, therefore, has NEP of 12.5 fW/√Hz and 15.7 fW/√Hz at 680-nm and 850-nm, respectively.

The SPADs have jitter of 113.2 ps as characterized by the FWHM of the measured IRF at 800 nm. The FW1/1000M (full width as measured at 1/1000 of the maximum value) is 8.05 ns.

#### VCSEL light sources

The 680-nm VCSEL (Vixar V00146) and 850-nm VCSEL (Light Avenue LM08NIV3) are chosen for their high optical bandwidth, generating optical pulses in response to electrical pulses down to 259-ps pulse width and helping to improve the temporal characteristics of the system IRF. Furthermore, in contrast to edge-emitting lasers, VCSELs require no additional optical components (such as prisms) to redirect light towards the tissue, which makes them favorable for dense integration in a wearable device. Mounting the VCSELs directly onto the surface of the ASIC and wirebonding the anode directly to on-chip driver electronics further improves integration density.

#### Control electronics on the chiplet

The chiplet includes active quench and reset circuits (AQC) for each of the 64 SPADs in the array, time-to-digital converters (TDC) with 65-ps resolution for timestamping photon detections, and a histogramming backend that supports collection of up to 1,048,575 photons per histogram bin. A digital controller allows for flexible configuration of integration time, VCSEL driver strength, and time-gating settings. Building on a previous chiplet design,^24^ this chiplet incorporates a dynamic reconfiguration controller, a shift register that accepts 17-bit packets that hold configuration settings such as SPAD time gating setting, VCSEL driver strength, VCSEL wavelength selection, and VCSEL enable. At any time, shifting in a new 17-bit packet causes the chiplet to modify its operation based on the newly requested settings. With this design, the FPGA can jointly control all 16 chips in the array, and can change which chiplet is actively emitting, which wavelength is active on that chiplet, and what the source and detector settings are during the source window, which is absolutely critical for being able to assemble complete frames of imaging data. Each chiplet consumes 60 mW during normal imaging operation. **Flexible PCB.** Micro-DOT is assembled on a flexible PCB, designed in Altium Designer and manufactured by Epec Engineered Technologies, that has a 50-µm polyimide core with only two metal layers to ensure that the device is flexible enough to be conformal to the tissue surface after assembly. 1/3 oz copper is used such that 3 mil trace and space is supported, which helps improve routing density and eliminate the need for extra metal layers. An electroless nickel electroless palladium immersion gold (ENEPIG) surface finish is used to prevent oxidation that could negatively affect flip-chip bonding yield. Matte black coverlay is used to cover the traces on the PCB, absorbing stray photons that could otherwise run along the surface of the PCB and contribute to background. A black selective soldermask, which can achieve much finer feature sizes than the coverlay, is used between adjacent flip-chip pads on the PCB to discourage shorting during flip-chip bonding. A polyimide stiffener is applied to the bottomside of the PCB where the PCB is inserted into the zero-insertion-force (ZIF) connector (Molex 541324062) on the motherboard to bring its total thickness to approximately 300 μm in that region. Laser cutouts in the polyimide expose the surface of each chiplet following flip-chip bonding.

Before beginning assembly of the PCB, the VCSEL cathode pads on the chiplets are gold-stud bumped and these bumped are “coined.” To perform this step, chiplets are mounted on glass slides (Fisher Scientific Premium Plain Glass Microscope Slides) using polymethyl methacralyte (PMMA) resist. The resist is spun on the glass slide and then cured on a hotplate (Thermo Fisher Scientific SP88854205) at 120C for 2-3 minutes. The cathode pads on the VCSELS are then stud bumped with gold bumps using a gold wire bonder (K&S IConn Plus). The gold bumps are then bump leveled using another glass slide with 15 N applied with a die bonder (Finetech Fineplacer Lambda). Chips are then released from the PMMA resist by submerging the glass slide in acetone overnight. Chips are removed from the acetone and washed with isopropyl alcohol (IPA), dried, and then stored for later use.

Assembly of the PCB begins by adhering the PCB to a four-inch silicon wafer (University Wafer 447) using polyimide film tape (3M 5413). Chiplets are flip-chip bonded to the PCB one at a time using a die bonder (Finetech Fineplacer Lambda) until all 16 chiplets are attached. Solder paste (Kester R276) is applied to pads on the PCB and decoupling capacitors are placed. The PCB and silicon wafer are then placed in a reflow over (Manncorp MC-301N) to complete attachment of the capacitors and also to ensure that flip-chip solder connections are uniform. After reflow, bump underfill is performed using a black, two-component underfill epoxy (Epo-Tek 353ND Black). The underfill epoxy is dispensed via a precision fluid dispenser (Nordson EFD Ultimus I) and then cured in an oven (Thelco 130D) at 150C for 5 minutes. Following underfill, the PCB can be safely detached from the silicon wafer.

The next assembly step is mounting of the VCSELs onto the surface of each chiplet. The PCB positioned to expose the surface of the chiplet through the cutouts on the PCB’s underside. A small amount of silver conductive epoxy (Epo-Tek H20E-D) is applied to both VCSEL cathode pads on every chiplet. A die bonder (Finetech Fineplacer Lambda) is used to pick-and-place each VCSEL onto the chiplet surface. A custom tool head for the die bonder is used to ensure that the VCSELs can be picked up without causing damage to the delicate topographical features on the VCSEL’s surface. Following VCSEL placement, the PCB is placed in a reflow oven (Manncorp MC-301N) at 120C for 15 minutes until the silver conductive epoxy has cured.

The anode of the VCSEL must be wire bonded to the VCSEL driver pad on the chiplet after mounting is complete. To provide stability to the flexible PCB during wire bonding, the PCB is attached to a custom holder that is 3D-printed using Clear Resin V4 (Formlabs) using a mounting adhesive (Crystalbond 555) and hotplate. The VCSEL’s anode pad is then wirebonded to the VCSEL driver pad on the chiplet. This bonding is performed at room temperature to avoid weakening the mounting adhesive. After wire bonding, the holder and PCB are separated by liquefying the adhesive at 80C on a hotplate. Residual mounting adhesive is removed in deionized (DI) water for 30 minutes.

Following VCSEL wire bonding, pinholes 3D-printed using a Biomed Black Resin (Formlabs) are mounted at each chiplet location using a moisture-cure, instant adhesive (Loctite 406). The 4-mm height of the pinhole far exceeds the height of the wire bond to the VCSEL anode, allowing the pinholes to also protect the wirebonds from damage during handling. The pinholes are filed down with a metal file prior to placement to ensure that they are smooth and flat on the top and bottom sides, and the height of each pinhole is identical.

In the final step of PCB assembly, a thermally conductive, room-temperature-cure flexible epoxy (Duralco 4538) is applied to the backside of each chiplet, which increases the surface area for heatsinking and also provides protection to the chiplet.

#### Interface motherboard

The interface motherboard is a rigid PCB designed in Altium Designer and manufactured by PCBWay. It contains all of the supporting electronics necessary to orchestrate the operation of the micro-DOT array. While size constraints for the motherboard are not as aggressive as they are for the flexible array, the motherboard actually contributes the majority of the total weight of the system (weighing approximately 93 g), and should still be kept as small as possible such that it can be easily mounted on the head. For this reason, we employ 4 layers on the motherboard with a 5 mil minimum trace width and spacing, which allows us to densely integrate discrete components and reduce overall motherboard size. We also reduce the thickness of the FR-4 substrate such that the total thickness of the board stackup is only 1 mm to further reduce unnecessary weight. The motherboard’s primary function is to create high-density connections between the FPGA module (Opal Kelly XEM7010) and the flexible array so that clocking and control signals can be distributed to the chiplets, and data outputs can be collected from the chiplets during imaging operation. The motherboard also hosts circuits that generate all required supply voltages from a single 5 V input. A buck converter (Analog Devices MAX20073) is used to generate the 1.5 V VDD rail, which is the core voltage for digital logic on the chiplet. A low dropout (LDO) regulator generates the 3.3 V HVDD rail for driving the VCSEL. A buck converter on the FPGA module is used to generate the 3.3 V VRST rail, which is used by the quenching circuits to reset the SPADs following an avalanche. A separate buck converter on the FPGA module is used to generate the 1.8 V DVDD rail, which sets the IO voltage of the chiplet. A boost converter (Analog Devices MAX5026) generates the 24.7 V SPAD rail, which sets the cathode voltage of the SPADs. Lastly, a Cuk converter (Texas Instruments LM2611) is used to generate a -5 V rail, which is paired with a digital potentiometer (Analog Devices MAX5436) and operation amplifier (Analog Devices LT1800) to provide a digitally-adjustable VCSEL cathode voltage. The motherboard has two 40-pin ZIF connectors (Molex 0541324062) that allow for direct connection of the flexible array. The motherboard is placed within a 3D-printed enclosure during imaging to isolate the subject from the electronics. The motherboard enclosure has an opening near the ZIF connectors to allow the flexible array to be connected, and another opening near the FPGA module for connection of power and data cables.

### System configuration for TD-DOT measurements

During HD-TD-DOT imaging experiments, the system is configured to operate at a specific frame rate for a specific illumination pattern. The most typical illumination pattern involves multiplexing between all 32 sources in the array and allocating each source an equal amount of integration time within the frame, with only a single source being active at any one time. For a 10-Hz frame rate, which is considered optimal for fNIRS^2^, this implies that each source is active for 3.125 ms (denoted as the “source window’) within the 100 ms spent collecting a single frame. The chiplets operate at 50 MHz, with the VCSEL being pulsed once each clock cycle, meaning that within a source window, the selected source is pulsed 156,250 times before moving on to the next source. While only a single source is enabled at any one time, all chiplets in the array actively collect photons and build ToF histograms during the source window. The chiplet can detect and timestamp as many as 8 photons per source pulse, meaning that a single chiplet is capable of collecting a maximum of 1.25×10^6^ total photons. At the end of each source window, all chiplets transmit their measured ToF histograms to the FPGA while simultaneously building a new ToF histogram that is associated with the next source. The FPGA packs the data from each source window into a frame and then stores the frame data in DRAM. Typically, after 10 frames have been collected, the data is shuttled over USB 2.0 to the PC. The system has a data rate of 1.92 megabytes per second during 10-Hz operation.

### Extracting features from ToF histograms

#### Integrated intensity

The integrated intensity parameter, E, is calculated as:

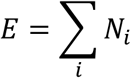

where Ni represents the number of photons in the i^th^ histogram bin. The summation of photons across time bins effectively eliminates the arrival time information, thereby transforming TD data into CW data.

#### Mean time

The mean time parameter, M, is calculated as:

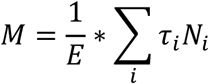

where τi represents the arrival time of the i^th^ histogram bin, Ni represents the number of photons in the i^th^ histogram bin, and E represents the integrated intensity of the ToF.

#### Laplace

The Laplace parameter, L, is calculated as:

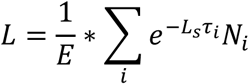

where τi represents the arrival time of the i^th^ histogram bin, Ni represents the number of photons in the i^th^ histogram bin, Ls is the Laplace coefficient, and E represents the integrated intensity of the ToF histogram.

Ls is chosen as −1 × 10^-4^ in this work because it provides high sensitivity for deeper layers while maintaining adequate SNR. In contrast, an Ls value of −1 × 10^-3^, for example, has less sensitivity but higher SNR, while an Ls value of −1 × 10^-5^ give excellent sensitivity but lower SNR (see **Fig. S12**).

### nEUROPt characterization

#### Phantom implementation

A tank having dimensions of 5 cm × 5 cm × 3 cm is 3D printed with Black Biomed Resin (Formlabs). The tank is open on the top, complete on all four sidewalls, and has small openings on the bottom that expose all of the chiplet locations in the array. The tank is filled with a solution that is composed of Intralipid, 20% emulsion (Sigma Aldrich) and India ink (Higgins) that are diluted in distilled water in specific ratios to achieve a modified scattering coefficient µ’s = 1.0 mm^-1^ and absorption coefficient µa = 0.01 mm^-1^. The absorption coefficient of the India Ink is measured prior to use in the phantom (**Fig. S15**). The cylindrical ROI having 5-mm diameter and 5-mm height is 3D printed from the same Black Biomed Resin (Formlabs) that is used for the tank. A small hole in the top of the ROI allows for a music wire (McMaster-Carr), painted with a flat white paint (Bic Wite-Out), to be inserted and glued in place. The music wire is attached to three linear translation stages (Thorlabs) that are stacked to allow for three axes of movement. In the nEUROPt protocol, the stepper motor controls the depth of the ROI relative to the SD plane.

#### Background and ROI captures

All nEUROPt results are presented as relative measurements, in which the difference it taken between a reference measurement taken of the imaging scene without the ROI (the background capture) and a separate measurement in the presence of the ROI (the ROI capture). For all nEUROPt results, the system was configured to collect ToF histograms with 20 ms of integration time for the selected source. 5000 total frames were collected in the case of both the background and ROI capture, making the total integration time per source 100 s.

#### Contrast

Contrast (C) is calculated considering all 100 s of integration time such that the effects of shot noise on the reported contrast values are minimal. The histograms from all 5000 frames of the background capture are averaged together into a single histogram, and then the integrated intensity of the resulting histograms is calculated and denoted as E_BKG_ . The same process is repeated for the ROI capture with the integrated intensity of the resulting histogram denoted as E_RO/_. Contrast is then calculated as:

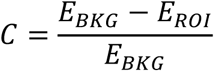

#### Contrast-to-noise ratio

Contrast-to-noise ratio (CNR) is calculated considering only 1 s of integration time such that the effects of shot noise are apparent in the result. The 5000 frames of the background capture are divided into 100 groups of 50 frames each. The histograms within each group are averaged together. The standard deviation of the integrated intensity across the 100 groups, C(E_BKG,1s_), is then calculated. E_BKG_ and E_RO/_ are calculated in the same way as is done for the contrast calculation, with all 5000 frames being included in the average. CNR is then calculated as:

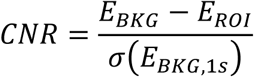

#### Mean time change

Mean time change (ΔM) is calculated considering all 100s of integration time such that the effects of shot noise on the reported mean time change values are minimal. The histograms from all 5000 frames of the background capture are averaged together into a single histogram, and then the mean time feature (M_BKG_) of the resulting histograms is calculated. The same process is repeated for the ROI capture to yield the mean time feature M_RO/_. Mean time change is then calculated as:

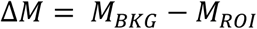

#### Mean-time-to-noise ratio

Mean-time-to-noise ratio (MNR) is calculated considering only 1 s of integration time such that the effects of shot noise are apparent in the result. The 5000 frames of the background capture are divided into 100 groups of 50 frames each. The histograms within each group are averaged together. The standard deviation of the mean time parameter across the 100 groups, C(M_BKG,1s_), is then calculated. M_BKG_ and M_RO/_ are calculated in the same way as is done for the mean time calculation, with all 5000 frames being included in the average. MNR is then calculated as:

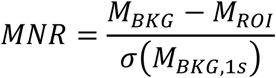

#### Laplace change

Laplace change (ΔL) is calculated considering all 100 s of integration time such that the effects of shot noise on the reported Laplace change values are minimal. The histograms from all 5000 frames of the background capture are averaged together into a single histogram, and then the Laplace parameter of the resulting histograms (L_BKG_) is calculated. The same process is repeated for the ROI capture with Laplace parameter of L_RO/_. Laplace change is then calculated as:

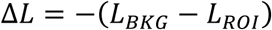

An inversion appears in the above equation such that Laplace change can be expressed as a positive number to be consistent with contrast and mean time change.

#### Laplace-to-noise ratio

Laplace-to-noise ratio (LNR) is calculated considering only 1 s of integration time such that the effects of shot noise are apparent in the result. The 5000 frames of the background capture are divided into 100 groups of 50 frames each. The histograms within each group are averaged together. The standard deviation of the Laplace parameter across the 100 groups, C(L_BKG,1s_), is then calculated. L_BKG_ and L_RO/_ are calculated in the same way as is done for the Laplace calculation, with all 5000 frames being included in the average. LNR is then calculated as:

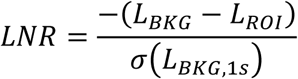

Just like for the calculation of the Laplace change, an inversion appears in the above equation such that the Laplace-to-noise ratio is a positive number to be consistent with contrast-to-noise ratio and mean-time-to-noise ratio.

#### Relative sensitivity

The relative sensitivity (S) of a given feature (*E*, *M*, or *L*) versus depth is calculated by normalizing that parameter to its peak value in the depth distribution. The sensitivity for E is given by:

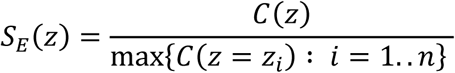

where *C* is contrast, and *n* is the number of depths for which contrast is measured. The sensitivity for *M* is given by:

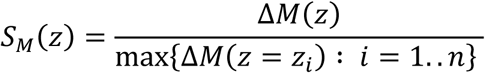

where ΔM is the mean time change and *n* is the number of depths for which mean time change is measured.

The sensitivity of *L* is given by:

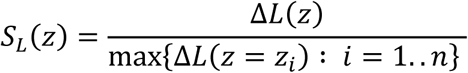

where ΔL is the Laplace change and n is the number of depths for which Laplace change is measured.

### DOT image reconstruction

#### Image reconstruction model

Tomographic reconstructions are generated using the PDE-constrained inverse model and the SENSOR (Sensitivity Equation-based Noniterative Sparse Optical Reconstruction) model^18^.

The PDE-constrained inverse model, which is known to be ∼30x faster than the existing nonlinear unconstrained optimization schemes, finds the unknown optical properties by solving the forward and inverse problems simultaneously within the framework of the following extended objective function called Lagrangian L(μ, u, η) as

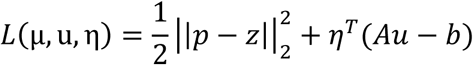

where u is the optical property vector, p is the prediction of the measurement z; Au − b is the forward model with the vector of light intensities u; and 7 is the vector of Lagrangian multipliers. The simultaneous solution of forward and inverse problems is achieved at a point where the gradient of L vanishes with respect to μ, u, and η.

The SENSOR inverse model, which has shown to be capable of performing approximately 1 mm^3^ spatial resolution optical tomographic imaging at a depth of ∼ 60 mean free paths (MFPs) in 20∼30 milliseconds, finds the unknown optical properties by solving the following regularized least squares problem:

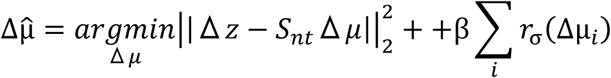

where Δz represents a differential measurement given in terms of logarithmic changes in the time-domain moments, E, M, or L, as Δz = Δln(E), Δln(M) or Δln(L); Δµ represents the differential change of target optical properties, such as µ_a_ and µ, or, in our case, [HbO2] and [HbR]; S is the non-truncated sensitivity matrix distinct from other linear approximations, f3 is a regularization parameter that balances between minimization of the data fit error term and the desired sparsity or smoothing level, and r_a_is the asymptotic ℓ_p_-norm function. Under appropriate assumptions^27,45^, especially for the retrieval of targets up to the depth of 2.5 cm^15,46^, our SENSOR model is simplified to an equivalent form of the linear model. The differential measurements and optical properties correspond to the change between the perturbed and unperturbed states; i.e. Δz = Δz _BKG_ − Δz _TARGET_ and Δµ = Δµ _BKG_ − Δµ _TARGET_, where the target could be the addition of an ROI in phantom studies or the beginning of a task or occlusion in the *in vivo* studies.

#### Chromophore recovery

Rather than computing the absorption values for each wavelength and then applying the Modified Beer-lambert law to calculate HbO2 and HbR, we directly calculate the chromophore concentrations using the spectral constraint technique, which has been shown to lead to smoother and more accurate reconstructions when compared to the two-step approach^47–49^. In this approach, measurements at each wavelength are coupled into one larger model, and the sensitivity matrix then relates the change in measurements to the change in chromophore concentrations rather than changes in absorption. The same PDE-constrained and SENSOR inverse models are then applied to solve for the unknown chromophore concentrations.

### Brain phantom experiments

#### Phantom implementation

A plastic shell having dimensions of 5 cm × 5 cm × 3 cm is 3D printed with Black Biomed Resin (Formlabs). The shell is open on the top, complete on all four sidewalls, and open on the bottom. The plastic shell is filled with three different layers of polydimethylsiloxane (PDMS, Dow Corning Sylgard 184). Scattering properties of the PDMS layers are controlled by mixing titanium dioxide (TiO2, Pantai Chemical USA PTR-620) into the curing agent prior to curing. Absorption properties of the PDMS layers are controlled by mixing India ink (Higgins) into the elastomer prior to curing. The ratio of elastomer to curing agent by weight is 10:1.

In the preparation, TiO2 is measured by weight and mixed into the curing agent with the resulting solution sonicated for 30 minutes to improve homogeneity. India ink is measured by volume and pipetted into the elastomer with the resulting mixture stirred for 10 minutes for homogeneity. The curing agent and the elastomer are mixed together and stirred for another 10 minutes. The uncured PDMS mixture is placed in a vacuum chamber (Sigma-Aldrich Scienceware vacuum desiccator) for 30 minutes to degas and remove any trapped air bubbles that could affect the optical properties of the PDMS layers.

The plastic shell is placed in a polytetrafluoroethylene (PTFE) evaporating dish (Fisher Scientific) with uncured PDMS mixture added to the desired volume. After placing in a vacuum chamber for 30 minutes to degas, the PDMS mixture is cured at 45C for 24 hours. This process is repeated two more times, with each subsequent layer being poured and cured directly atop the previous layer, until a total of three PDMS layers (representing the optical properties of scalp, skull, and CSF) are fabricated.

The cured phantom (**Fig. S16**) is then filled with a solution that is composed of Intralipid, 20% emulsion (Sigma Aldrich) and India ink (Higgins) that are diluted in distilled water in specific ratios to achieve the desired optical properties of the gray matter. A cylindrical ROI is 3D printed from Black Biomed Resin (Formlabs). The size of the ROI is either 5 mm diameter and 5 mm height, or 3 mm diameter and 3 mm height depending on the experiment. The ROI is suspended with music wire (McMaster-Carr) that is painted with a flat white paint (Bic Wite-Out). The music wire is attached to three linear translation stages (Thorlabs) that are stacked to allow for three axes of movement. In the brain phantom experiments, the stepper motor controls the depth of the ROI relative to the SD plane, as well as the lateral XY position of the ROI relative to the array.

#### Background and ROI captures

Brain phantom measurements are presented as relative measurements, as is done in the nEUROPt experiments. A reference measurement is taken of the imaging scene without an ROI (the background capture), and a separate measurement is taken where the ROI is included in the imaging scene (the ROI capture). Differential changes between the background and ROI capture are used as the basis for image reconstruction. A single background capture is sufficient for the reconstruction of all ROI locations. The system was configured to collect ToF histograms with 20 ms of integration time for the selected source. 50 total frames were collected in the case of both the background and ROI capture, making the total integration time per source 1 s.

#### Image Reconstruction

Prior to image reconstruction, all 50 of the imaging frames in the background capture are averaged together to create a single representative frame. In this frame, each source window has 1 s of integration time. The same is done for the ROI capture, and then the differential changes between the background and ROI capture are used for image reconstruction.

#### Localization error

Localization error is a metric used to characterize the error in correctly identifying the location of the centroid of the ROI in the final reconstructed image. It is represented by the red vector in the reconstructed images of **Fig. 4C**. A threshold is first applied to the reconstructed image to produce a binary image with a reconstructed target of uniform intensity. The x-coordinate and y-coordinate of the centroid of the reconstructed target (denoted xc,rb and yc,rb, respectively) are found by calculating the x- and y-moment of the image. The localization error is calculated as the length of the vector pointing from the centroid of the reconstructed target to the centroid of the ground truth object location, which has x-coordinate (*xc,gt)* and y-coordinate (*yc,gt*):

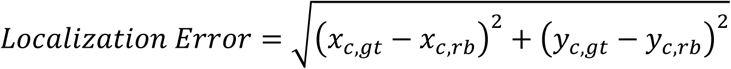

#### Effective Resolution

Effective resolution is a metric used to characterize both the localization error and the error in correctly identifying the size of the ROI in the final reconstructed image. It is represented by the yellow circles in the reconstructed images of **Fig. 4C**. A threshold is first applied to the reconstructed image to produce a binary image with a reconstructed target of uniform intensity. The distance between the centroid of the ground truth object location and each of the pixels in the reconstructed target is calculated. The largest of these distances defines the radius of a circle which, when drawn to be centered on the centroid of the ground-truth object location, completely encloses the reconstructed target. The diameter of this circle represents the effective resolution.

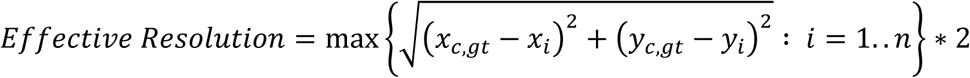

where n is the number of pixels in the reconstructed target, and a given pixel has x-coordinate xi and y-coordinate yi.

### Forearm imaging

#### Experimental setup

The experiment begins with the subject seated comfortably in a chair with their right hand resting on a desk. A blood pressure cuff (BPC, Dixie EMS) was positioned just above the elbow and inflated to a pressure of 60 mmHg. A visible vein on the underside of the forearm was identified as the location for positioning the Micro-DOT array during the experiment. The subject’s hand was positioned under the motherboard, palm facing upward, to facilitate the placement of the array on the forearm’s underside. A cinch strap (Envisioned Velour) was securely tightened around the array, ensuring good coupling throughout the procedure. During the occlusion phase, the BPC was rapidly inflated to 60 mmHg, and this pressure was maintained as consistently as possible to ensure reliable experimental conditions. At the end of the occlusion phase, the BPC was rapidly deflated. Care was taken during BPC inflation and deflation to ensure that the BPC did not cause motion artifacts. The Micro-DOT system was configured to image with a frame rate of 5 Hz, implying an integration time of 6.25 ms per source.

#### Image reconstruction

Prior to reconstruction, a high-pass filter with a cutoff of 0.01 Hz was applied to the raw measurement data to correct for baseline wander or DC offset. Furthermore, as the expected hemodynamic change occurs over a period of several seconds, we downsampled the data from 5 Hz to 1 Hz to reduce the number of reconstruction tasks. The first 10 frames of the baseline measurement period were averaged to use as the representative background value in the differential measurements for the image reconstruction algorithm.

### Forehead imaging

#### Experimental setup

The breath-hold experiment is designed with the subject seated in an upright position with the Micro-DOT device positioned on the forehead, attached with a cinch strap. The subject is instructed to remain as still as possible during the inhalation and exhalation surrounding the breath-hold to minimize motion artifacts. The Micro-DOT system is configured to image with a frame rate of 5 Hz, implying an integration time of 6.25 ms per source.

#### Image reconstruction

Raw experimental data is preprocessed using a Gaussian moving average filter with a window size of 40 to reduce high-frequency noise components. After that, the data is downsampled from 5 Hz to 1 Hz, and the first 60 frames (1 minute) are averaged to represent the baseline measurement for differential measurement-based reconstruction. The following 120 frames are then used for reconstruction, covering a 40-second breath hold and 1-minute recovery.

### Brain imaging of motor cortex with finger tapping

#### fMRI scans

Placement of the Micro-DOT device during finger tapping experiments is guided by structural and functional fMRI scans performed prior, wherein the subject is asked to follow the same finger tapping protocol that will be used in the Micro-DOT experiment that will follow. Imaging was acquired on a Siemens Prisma with a 64-channel head coil. A T1w 3D MP-RAGE structural scan was collected with a repetition time (TR) of 1900 ms, a flip angle of 9 degrees, an echo time (TE) of 2.52 ms, voxel dimensions of 0.976562 mm × 0.976562 mm × 1 mm, and matrix dimensions of 256 × 256 ×176. Four functional EPI runs were collected with a TR of 1200 ms, a flip angle of 65 degrees, a TE of 30 ms, voxel dimensions of 2 mm × 2 mm × 2 mm, and matrix dimensions of 112 × 112 × 66. A total of 234 time points were collected.

The fMRI data collected during the experiment was preprocessed using default parameters in FSL’s FEAT Version 6.0, including motion correction, slice timing correction, spatial and temporal filtering. A GLM analysis was performed in which the task regressor representing the finger tapping blocks is convolved with the individual subject’s double-gamma hemodynamic impulse response function (HRF). The HRF was estimated from an independent experiment as described in Grinband et. al^50^. A fixed-effects group analysis was performed across runs. The z-statistic image is thresholded at z=3.1 with a cluster threshold of p < 0.05 using Gaussian Random Field theory, binarized, and intersected with a mask of the pre-central gyrus to create the motor cortex region of interest.

#### Experimental setup

The experiment begins with the subject sitting upright in a chair. The structural scan and functional overlay MRI scans for the subject are loaded in Brainsight (Rogue Research). A passive rigid body is attached to the subject’s head with a small adhesive patch and is tracked by the Brainsight system. Registration of the subject’s position is performed using a pointer that is touched against predetermined location on the subject’s head and tracked by the Brainsight system. Following registration, we determine the appropriate location for the Micro-DOT array by finding the location on the scalp directly above the activated region shown in the functional overlay. A skin-safe marker is used to make a yellow mark at the corresponding location on the scalp. The Brainsight system is then switched off to prevent the flashing of the Brainsight system’s light source from creating artifacts in the Micro-DOT data. The device is placed over the yellow mark and secured with a 3.5-inch-by-4-inch wound-dressing (3M Tegaderm). A cinch strap is secured across the patient’s head, the motherboard is connected to the Micro-DOT array, and then the motherboard is secured to the cinch strap using hook and loop fasteners. Additional cable straps are added that run across the Micro-DOT array to give better coupling.

#### Scalp coupling index (SCI) characterization

To assess the coupling of SD locations to the scalp after device placement, we calculate the SCI at each for each channel. For a given channel, the SCI is calculated by first transforming the measurement to CW by considering only *E* for both wavelengths and bandpass filtering the resulting signal between 0.5 Hz and 2.5 Hz such that the dominant component of the filtered signal is contributed by the heart rate. The filtered signals are divided by their standard deviation and then the cross-correlation is computed between the two wavelengths. The *E* data produced by our TD device tends to be noiser than equivalent data generated by dedicated CW devices because the VCSEL is duty cycled for TD operation, resulting in a significant reduction in average optical power. This means that the value of the SCI is not just affected by the coupling of the SD to the tissue, but also by the SDS of the channel and SNR of the measurement. For this reason, we typically assess coupling by looking at the maximum SCI for each detector considering those sources that immediately surround it, which eliminates false negatives caused by low SNR at large SDS.

When conducting an experiment, the Micro-DOT device is placed on the scalp above the target brain region, and then about 30 s of frame data is collected with the subject entirely at rest. The SCI values across the Micro-DOT device are then calculated to ensure that the coupling across the device is adequate and relatively uniform (**Fig. S3B**). If the SCI figure reveals that a corner of the device is poorly coupled, the device can be repositioned prior to running the actual imaging experiment.

#### GLM analysis

GLM is a well-known technique for analyzing CW fNIRS data, and toolboxes such as Homer3 and NIRS Brain AnalyzIR Toolbox have made analysis of CW fNIRS data very accessible and helped to standardize the analysis pipeline. The HD-TD-DOT data from Micro-DOT was reduced to CW-DOT data by summing photons in the ToF histograms and removing channels with intermediate SDSs, which we expect will not be informative when only CW information is considered. The resulting dataset was processed using the NIRS Brain AnalyzIR Toolbox. It is more difficult to achieve statistical significance in GLM using the CW information from a TD-DOT device, because the intentional use of a pulsed source results in a significant reduction in average power (∼25× in our case) through duty cycling, reducing SNRs. To compensate for this reduction, we use only a single source location and devote all integration time within each frame to that source location, resulting in a 16× increase in integration time.

#### FLOBS fitting

Temporal fitting is performed with three basis functions taken from FMRIB’s FLOBS. The six-second duration boxcars of the finger tapping task were convolved with the basis functions. Following the convolution, a non-linear least squares fitting function (SciPy’s curve-fit function) is used to find the coefficients of each basis function that minimized the error between the predicted basis function and the measured hemodynamic response. The [HbR] time trace is inverted with the expectation that decreases in concentration of [HbR] are indicative of increases in activation. An illustration of the convolved basis functions and fit coefficients is shown in **Fig. S17**. The HRF was created by taking a weighted mean of the original basis functions, weighted by the fit coefficients. The [HbO2] and [HbR] HRFs can then be compared directly the BOLD HRF extracted using fMRI.

#### Micro-DOT image reconstruction

Prior to image reconstruction, raw experimental data are filtered and then block averaged across the 30 finger tapping trials. Filtering was performed on raw ToF histograms rather than extracting E, M, and L features beforehand and filtering afterwards. A time trace for each ToF histogram bin over the course of the 30 trials is filtered using a fifth-order digital Butterworth filter with a cutoff frequency of 0.3 Hz. The filter is applied forward and backward to ensure that it did not introduce delay in the filtered signal, which ensured that the timing of the data remained consistent with the onset of the finger tapping blocks. Block averaging is then performed by windowing the time trace of each ToF histogram bin between 2-s before and 20-s after the onset of each finger tapping block. Time traces in each window are averaged together, with the result effectively representing a block averaged and filtered version of the ToF histograms that are collected in response to all 30 finger tapping tasks.

The E, M, and L features are extracted from the resulting block-averaged ToF histograms. The time traces of these extracted features are then processed using a Gaussian moving average filter with a window size of 10. Additionally, these time traces are further filtered using a fifth-order digital Butterworth low-pass filter, which is to eliminate task-unrelated systemic physiological signals, such as heart rate which typically oscillates around 1 Hz, fluctuations in blood flow and oxygenation levels due to respiratory cycles (approximately 0.2 Hz), and Meyer waves with low-frequency oscillation in venous blood flow (around 0.1 Hz). Long-channel time traces, corresponding to brain activity and characterized by a low signal-to-noise ratio (SNR), are filtered using an aggressive low-pass filter with a cutoff frequency of 0.09 Hz. In contrast, short-channel time traces, which reflect scalp activity and have a high SNR, are processed with a low-pass filter that has a cutoff frequency of 0.3 Hz.

For the baseline measurement, the first ten frames (2 seconds) prior to the onset of finger tapping are taken, filtered with a low-pass filter that has a cutoff frequency of 0.009 Hz, and averaged to represent the steady baseline value. This baseline value is used as a reference point against which each frame is compared to calculate the differential changes in [HbO2] and [HbR], leading to the hemodynamic response over a time window spanning from 2 seconds before to 14 seconds after the finger tapping block. Reconstructions of the differential changes of HbO2 and Hb concentrations were performed with the SENSOR inverse model as described earlier using both short- and long-channel temporal data. Depth-dependent regularization^51,52^ was also applied in the SENSOR reconstruction to differentiate between scalp and brain activities at different depths effectively. This depth-dependent regularization improves the conditioning of the sensitivity matrix in the SENSOR algorithm by providing higher weights to the regions with lower sensitivity. Specifically, stronger regularization was applied to deeper depths such as the brain region where sensitivity is lower, and weaker regularization was used to swallow depths such as the scalp region, where sensitivity is very high. As a result, depth-dependent regularization provides a more uniform spatial resolution and contrast across the entire field-of-view (FOV) imaging domain.

#### Overlay process

In order to perform overlap of Micro-DOT images with fMRI, there are several steps that must be taken. First, the reconstruction volume must be registered at the correct position in the T2 scan, and then the reconstruction volume must be resampled and interpolated such that it has equivalent volume and voxel size to the T2 scan. In Python, we create a mask of the reconstruction volume and insert alignment points at the physical corners of the Micro-DOT array. In FSL’s FLSEyes tool, we then create a mask of the T2 volume, and alignment points corresponding to the physical corners of the Micro-DOT array are added to the volume based on the placement of the array during the experiment. FSL’s FLIRT tool is configured to register the mask of the reconstruction volume against the mask of the T2, both being empty with the exception of the alignment points corresponding to the corners of the Micro-DOT array (**Fig. S18**). A mutual information cost function is used, and the reconstruction volume mask is treated as a rigid body meaning that only 6 degrees of freedom (x position, y position, z position, pitch, roll, and yaw) are used for registration. The resulting transform is then applied to the true reconstruction volume such that it is mapped into the T2 volume (**Fig. S19**).

Once the reconstruction volume has been mapped into T2 space, brain extraction is performed using FSL’s BET tool. Standard brain extraction using bet2 was performed with the T2 as an input image, and a fractional intensity threshold of 0.35 with a threshold gradient of -0.15. A second standard brain extraction using bet2 was performed with the previous brain extraction result as an input image, and a fractional intensity threshold of 0.35 and a threshold gradient of -0.15.

The resulting T2 brain volume coming out of BET is then transformed into MNI space using FLIRT (**Fig. S20**). MNI152_T1_2mm_brain was used as a reference image for the transformation. 12 degrees of freedom are used with a mutual information cost function. The resulting transformation is applied to the Micro-DOT reconstruction volumes as well as the functional data from MRI (**Fig. S21**).

The next step in the overlay process is to perform volume-to-surface mapping of the reconstruction volumes onto the left hemisphere. This is done using Connectome Workbench’s volume-to-surface-mapping tool with reference surface Q1-Q6_R440.L.midthickness.32k_fs_LR.surf.gii. To understand what the FoV of the device looks like at the end of the process, we create a sample reconstruction volume of all 1s and carry it through this process. The result is a precise definition of which regions of the brain the device is capable of measuring activity given its position on the scalp (**Fig. S22**).

We present surface heatmaps of Micro-DOT data that are derived from HbO2 and HbR chromophore concentration changes within the FoV. To create the surface, we first take the HbO2 and HbR concentration volumes and perform volume-to-surface-mapping to reference surface Q1-Q6_R440.L.midthickness.32k_fs_LR.surf.gii. The HbO2 and HbR surfaces are added together to create an HbT surface, which is used in the surfaces presented in **Fig. 6K**. HbO2 and HbR surfaces are shown in **Fig. S23**.

For comparison of the localization capabilities of the Micro-DOT device with fMRI, we create new surfaces that show only strongly activated regions by assigning value 1 to all pixels with amplitudes in the 75^th^ percentile or higher, and all other pixels in the 74^th^ percentile or below value 0. Because the FoV was primarily activated pixels as opposed to deactivated pixels, and because the activation was of much higher amplitude than the deactivation, we focused only on the comparison of the location of strongly activated regions and did not compare the location of strongly deactivated regions. Outlines of the strongly activated regions detected using fMRI were overlaid on top of the strongly activated regions detected using Micro-DOT. fMRI volume data is masked using the FoV volume (the same volume that is used to create the surface in **Fig. S22)**. All surface data is shown on reference surface Q6_R440.L.inflated.32k_fs_LR.surf.gii so that the FoV can be better visualized.

#### Dice Coefficient

The Dice coefficient is a metric for quantitatively characterizing the agreement between two surface heatmaps, in this case, the fMRI and Micro-DOT cortical surface heatmaps.

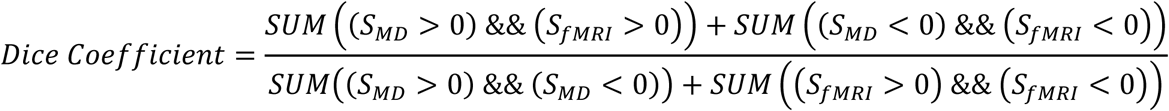

where SMD is the Micro-DOT surface and SfMRI is the fMRI surface. In this case, the greater than and less than operators are evaluated at each pixel on the surface, and return a new surface with 1s in those pixels where the logical operation is true, and 0s in those pixels where the logical operation is false. The && (logical AND) operator is also a logical operator that is evaluated at each pixel on the surface and returns a new surface with 1s in those pixels where the logical operation is true, and 0s in those pixels where the logical operation is false. The SUM operator is a reduction operator, which sums the values of all pixels on the surface and returns a single value.

#### Study Protocol Approval and Informed Consent

The study protocols for all experiments making use of human participants were approved by Columbia University’s Human Research Protection Office and Columbia’s IRB regulations under approved IRB protocol AAAT4908 “Use of a time-domain diffuse optical tomography (TD-DOT) imaging array for brain imaging in the near infrared (NIR).” Informed consent was received from all participants. Consent to publish identifiable images was obtained from relevant participants.

