## Supplementary Information for "Wearable, High-Density, Time-Domain Diffuse Optical Tomography Array for Functional Neuroimaging"

### Section S1: SPAD characterization of design-of-experiment arrays

To determine the best SPAD structures to use in Micro-DOT, we fabricated device arrays for a design of experiments (DOE) and built automated test setups to extract current-voltage (IV) curves, to perform DCR/PDP characterization, and to measure the IRF of 32 different SPAD structures. Variations in the SPAD structures that were tested in the DOE array are shown in **Fig. S24** and **Fig. S25**. Characterization results for all SPADs that were fully tested is presented in **Table S3**.

**Current-voltage measurement.** The purpose of the IV measurement setup (**Fig. S26**) is to verify that the SPAD shows appropriate diode characteristics and to determine the breakdown voltage. The setup includes a motherboard that hosts an Arduino, that is responsible for controlling drivers that enable and disable 64 mechanical relays, and BNC connectors that allow a Keithley 2400 Sourcemeter to be connected to the anode, cathode, and substrate rails on the motherboard. Relays are used instead of transistors because of their negligible leakage current. The system also includes a daughterboard that is connected with two 40-pin, zero-insertion force (ZIF) cables to the motherboard and hosts a 100-pin quad flat no-lead (QFN) package containing the SPAD array chip. The daughterboard also includes quenching resistors for each SPAD, which are connected with a 0.1” shunt but which are disconnected during IV measurements.

During operation, a Python program is used to jointly control the Keithley and the Arduino. The Arduino is controlled with a serial interface; drivers buffer the digital outputs of the Arduino to drive reasonably high currents (~50 mA) to control the relays. Two relays are enabled at any one time and electrically connect a single SPAD’s anode and cathode terminals to the Keithley. The cathode voltage is swept by reconfiguring the Keithley’s output voltage via the Python program. Each time that the output voltage is changed, a configurable number of current readings are collected from the Keithley and averaged together to create a point on the IV curve. During the sweep, the SPAD is exposed to ambient light to ensure that it breaks down as soon as the breakdown voltage is exceeded. This process is repeated until a full IV curve has been created. It is important to also note that the Keithley is configured with a compliance current limit to ensure that the structures are not damaged by large currents flowing during avalanche breakdown. A sample IV curve result from a single SPAD is displayed in **Fig. S27**.

**PDP and DCR characterization.** A second test setup (**Fig. S28**) was used to characterize the PDP and DCR of the SPADs in the same DOE array. In this case, the motherboard hosts an XEM7360 FPGA module (Opal Kelly) for digital control, 64 mechanical relays for electrically connecting the anode and cathode of a single SPAD in the DOE array, drivers for buffering digital outputs of the FPGA to control the relays, a fast comparator for digitizing SPAD avalanche currents, BNC connectors for connection of a Keithley 2400 Sourcemeter, and two 40-pin ZIF connectors for connection to the daughterboard. The daughterboard is the same as that used for the IV curve test setup, but in this case, the 0.1" shunt is connected such that 100-kΩ quenching resistors are connected between the anode of each SPAD and ground.

For PDP measurement, a Xenon fiber optic light source (Spectral Products, Model Number Y1603, 175W Ozone-blocking) is used to create broadband light across a large portion of the visible and infrared spectrum. A monochromator (Spectral Products, Model Number CM110) is used to filter this white light with 0.6mm slits to set the linewidth at 5 nm. The output of the monochromator is fed via optical fiber into one port of an integrating sphere (Newport, Model Number 819C). At another port of the integrating sphere, a reference photodiode (Newport, Model Number 818-IS-1) is connected, and the output of this photodiode is connected to a benchtop powermeter (Newport, Model Number 1936-R). At another port of the integrating sphere, the daughterboard is mounted vertically such that the entire QFN package sits within the integrating sphere (see **Fig. S28**).

A Python program automatse the DCR measurement for each SPAD in the DOE array. The FPGA is configured to enable the relays on the motherboard such that a single SPAD's cathode and anode are connected. The Keithley is configured to set the cathode voltage of the SPAD. The FPGA enables a counter that takes input from the comparator on the motherboard and begins counting clock cycles to guarantee a specific measurement time (configurable in Python, typically 10s). SPAD avalanches are digitized by the comparator and then captured by the counter on the FPGA. At the conclusion of the 10-s measurement window, the Python program reads the number of detected avalanches from the FPGA. This process is repeated for different excess bias voltages, and then the whole process is repeated for all 32 SPAD structures on the DOE array.

For PDP measurement, the FPGA is configured to enable the proper set of relays such that a single SPAD's cathode and anode are connected with the Keithley configured to set the cathode voltage of the SPAD under test. The center frequency of the monochromator is set, and the FPGA enables its counter and begins counting clock cycles exactly as is done in the DCR measurement. At the same time, optical power readings are collected from the powermeter. At the conclusion of the measurement, the Python program collects the number of photons from the FPGA and averages the optical power reported by the powermeter. This process is repeated for many different monochromator wavelengths, many different excess bias voltages, and for all of the structures in the DOE array. A sample PDP plot is shown in **Fig. S14B**.

The integrating sphere ensures that the light intensity at the daughterboard port and the reference photodiode port is the same. By comparing this number of photons per unit time to the number of photons detected by the SPAD per a unit time, the photon detection probability can be calculated as:

$$PDP\ (\%) = \frac{\left(\frac{N_{SPAD}}{t_{meas}} - DCR\right) * E_{photon}}{A_{SPAD}} * \frac{A_{PD}}{P_{PD}} * 100$$

where $N_{SPAD}$ is the number of photons collected by the SPAD during the measurement time $t_{meas}$, DCR is the dark count rate for the SPAD in Hz, $E_{photon}$ is the energy of the photon at the measurement wavelength, $A_{SPAD}$ is the photosensitive area of the SPAD structure, $A_{PD}$ is the photosensitive area of the reference photodiode, and $P_{PD}$ is the power measured by the reference photodiode. The DCR is subtracted from the number of photons measured by the SPAD.

**Impulse response function (IRF) characterization.** The IRF measurement setup (**Fig. S29**), unlike the IV curve and DCR/PDP setup, was not automated due to the complexity of doing so while preserving the high-bandwidth nature of the measurement. For this reason, only selected SPADs that have adequate DCR and PDP are carried forward to IRF testing. The DOE array die is attached to a small PCB that allows for bonding of up to eight SPADs. Only a small number of SPADs are bonded out to ensure that wire bond length is minimized and the bandwidth of the measurement is not adversely affected by parasitic inductance. The PCB also has spaces for small quenching resistor, and pads that allow for probing of the anode voltage of the SPAD. A 3.5-GHz differential active oscilloscope probe (Tektronix P7330) is connected across the quenching resistor for the IRF measurement.

A tunable, femtosecond Ti:Sapphire laser (Coherent Chameleon Vision II) is used as a pulsed light source. The optical power of the laser is attenuated by adjusting the coupling of the laser into an optical fiber such that the SPAD sees one to five photons for every 100 laser pulses, thereby mitigating distortion of the IRF due to photon pileup. To provide a timing reference for incoming photons, which is necessary for an IRF measurement, the output of the laser is split between the main optical path, which runs to the SPAD, and another optical path that runs to a photodiode (Thorlabs S120C). The output of the photodiode creates an electrical pulse in response to the optical pulse, providing a timing reference. Because the photodiode can only measure wavelengths at or above 800 nm, all IRF measurements were performed at this wavelength.

The photodiode output as well as the output of the differential probe are fed to a 23-GHz oscilloscope (Tektronix MSO 72304DX) which has a built-in function for measuring timing jitter between a reference clock signal (from the photodiode) and a data signal (from the differential probe). The oscilloscope is configured for a time-interval error (TIE) jitter measurement, which calculates the delay between the reference clock and the data signal over many clock cycles and creates a histogram from the results. A representative IRF measurement is shown in **Fig. S14C**.

Characterizing the full width at 1/1000 of the maximum (FW1/1000M) of the IRF is more challenging because the measurement typically does not contain enough photons to decay three orders of magnitude from the peak before hitting the noise floor created by the DCR. As a result, some extrapolation is used to extract an approximate FW1/1000M. Well past the peak, the decay of the IRF is purely exponential and driven by the diffusion of photogenerated carriers towards the multiplication region[1]. The time constant of this decaying exponential is given by

$$T = \frac{W^2}{\pi^2 D_n}$$

where W is the thickness of the plane neutral region and $D_n$ is the minority carrier diffusion coefficient. The minority carrier diffusion coefficient is given by

$$D_n = \mu_n \cdot \frac{kT}{q}$$

where $\mu_n$ is the minority carrier mobility, k is Boltzmann's constant, T is temperature, and q is the minority carrier charge. Given this purely exponential decay, we can confidently fit the tail of the normalized IRF with a decaying exponential function and extrapolate down to $10^{-3}$ to find the FWHM. This process is illustrated in **Fig. S14D**.

### Section S2: Extended GLM Discussion

#### Finger Tapping Experiment – 6-Second Duration

GLM results show statistically significant ($p < 0.05$) [$HbO_2$] response in the measurement channel most directly overlapping the region of expected activation, and statistically non-significant ($p \geq 0.05$) activation elsewhere (**Fig. S30A**). We did not find statistical significance for [HbR] in any of our channels for this experiment (**Fig. S30B**). It is clear from the block averages (**Fig. 6D**) that all channels, even those with larger SDS, have a temporal response that is dominated by task-evoked scalp activation. For a 6-s finger-tapping task, we would expect that the [$HbO_2$] signal originating from the motor cortex would rise following the onset of the task, and then remain at a level higher than the baseline until sometime after the task block has concluded. In contrast to this, it can be observed in our block average data that the [$HbO_2$] signal begins to decay back towards the baseline value before the 6-s finger tapping task ends. This is likely because the dominant component of the signal originates from task-evoked $HbO_2$ activity in the scalp.

As shown in **Fig. 6E**, one of the steps in the GLM pipeline is regression of short channel principal components. This, in theory, should remove task-evoked signals originating from the scalp, leaving behind only those signal components originating from the motor cortex. However, even after this step, the temporal profile of the remaining signal still decays earlier than the end of the 6 s finger tapping task (**Fig. S31**). This negatively affects the statistical significance of the GLM fit results.

#### Finger Tapping Experiment – 10-Second Duration

We also performed a finger tapping experiment with a 10-second duration that was in every other way identical to the 6-second duration finger tapping experiment (**Fig. S32A**). Brainsight was used to place the array on the head directly over the region of the brain that we expected to be activated by the 10-s finger tapping task (**Fig. S32B**). We once again calculated our SCI values prior to performing the experiment (**Fig. S32C**) to verify coupling of the device to the head. For the 10-second experiment, the array that was used had many sources that were not properly working, as well as a few detectors that were not functional. As a result, spatial reconstructions from the 10-second experiment were not possible, but GLM analysis was still possible for the channels that were functional. GLM results showed statistically significant $HbO_2$ activity for several of the channels overlapping with the expected ROI. Relative to the 6-s finger tapping experiment, the difference in the temporal dynamics of channels between short and long SDS is much more apparent in the 10-s finger tapping experiment. Channels with short SDS show a very rapid increase in $[HbO_2]$ in response to the onset of the task, but $[HbO_2]$ fall sharply well before the 10-s finger tapping task is completed (see **Fig. S32D**, upper left plot). In contrast to this, channels with longer SDS have very different temporal dynamics, with a sharp increase in $[HbO_2]$ occurring following the onset of the task and persisting through the end of the 10-s finger tapping task (see **Fig. S32D**, lower right plot). For the 10-s finger tapping experiment, we saw more statistical significance in our channels despite the fact that we performed only 10 repetitions of the 10-s finger tapping task compared to 30 repetitions of the 6-s finger tapping task (see **Fig. S32F**), likely due to the better temporal separation of the task-evoked scalp and brain signals that comes with a longer task duration.

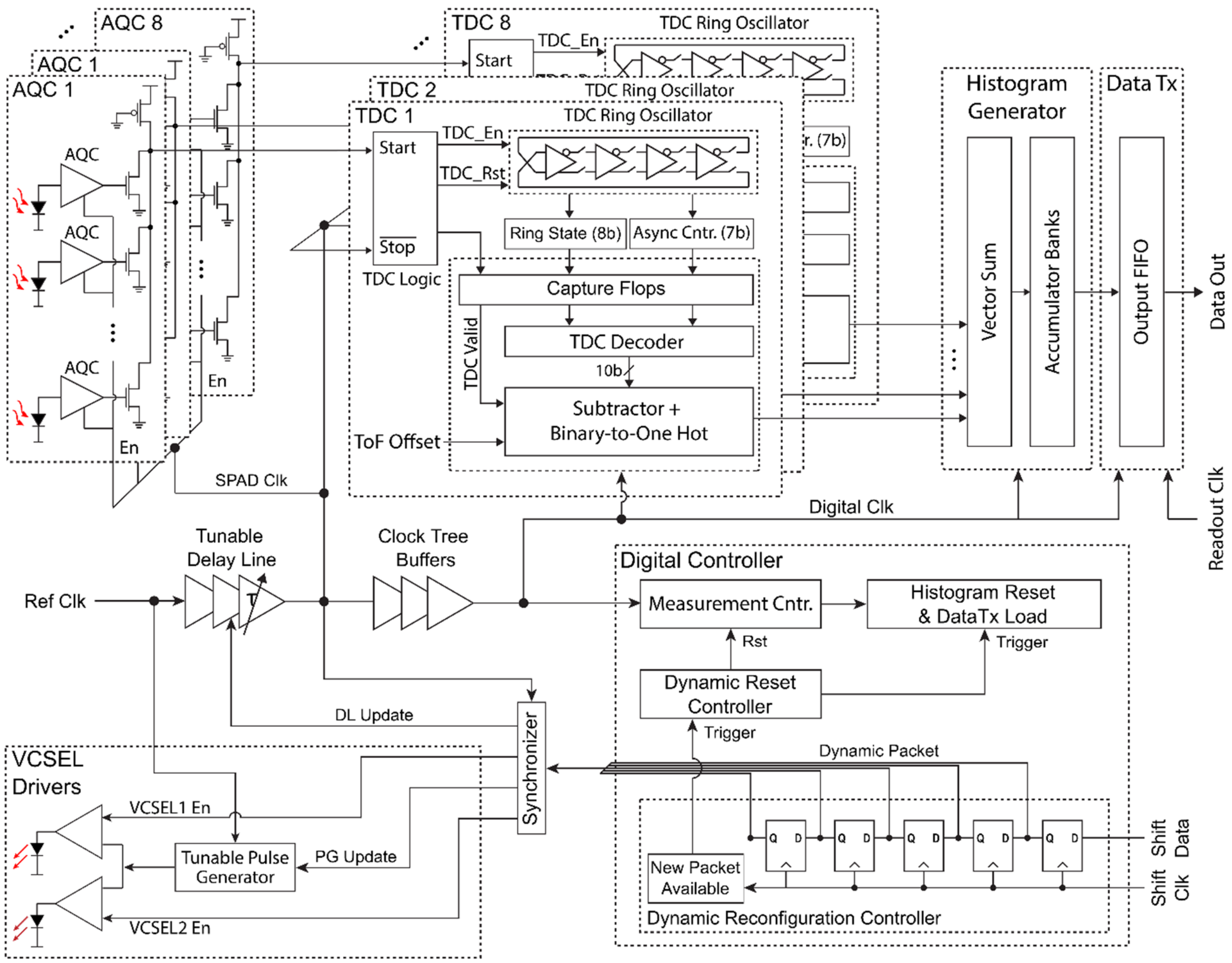


**Fig. S1. Micro-DOT chiplet block diagram.** The important hardware blocks, including the AQC rows, TDCs, VCSEL drivers, and digital controller are shown. Specific detail is given to the distribution of the reference clock into three separate domains that clock the VCSEL drivers, TDCs/SPADs, and digital controller separately. The dynamic reconfiguration controller is highlighted for its relevance to this work. This controller allows for on-the-fly reconfiguration of the chiplet, including whether the VCSEL is enabled, which VCSEL wavelength(s) is active, VCSEL driver strength, and SPAD time gating delay. In the system, this controller is the main mechanism through which the FPGA can orchestrate multiplexing between different sources on the flexible array to create a frame.

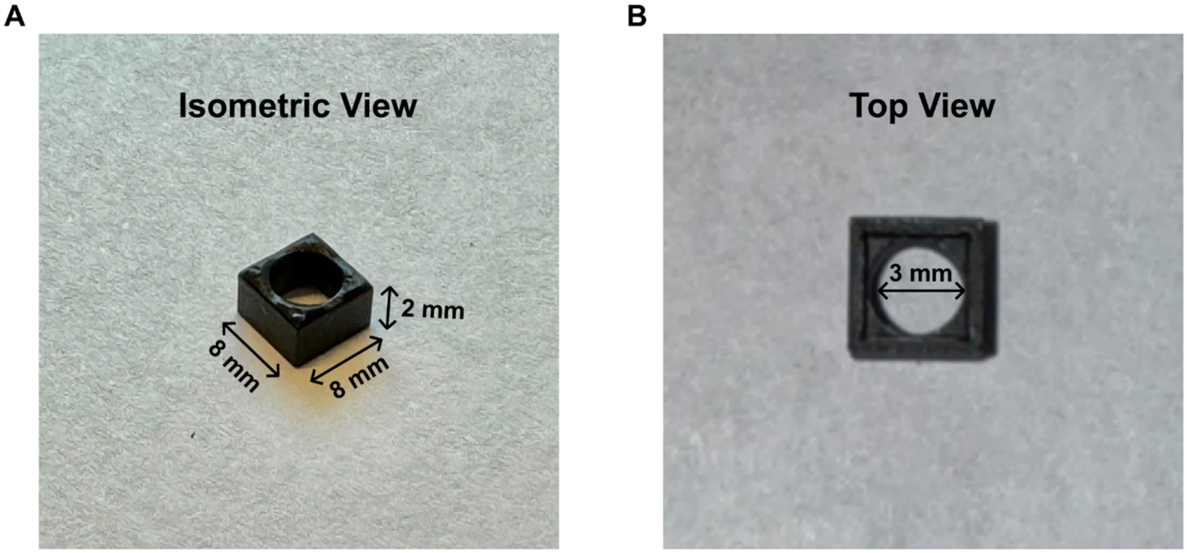


**Fig. S2. Pinhole structure mounted at each SD location.** (A) Isometric view of pinhole showing outer dimensions of the square pinhole. (B) Top view of pinhole showing inner diameter of circular aperture.

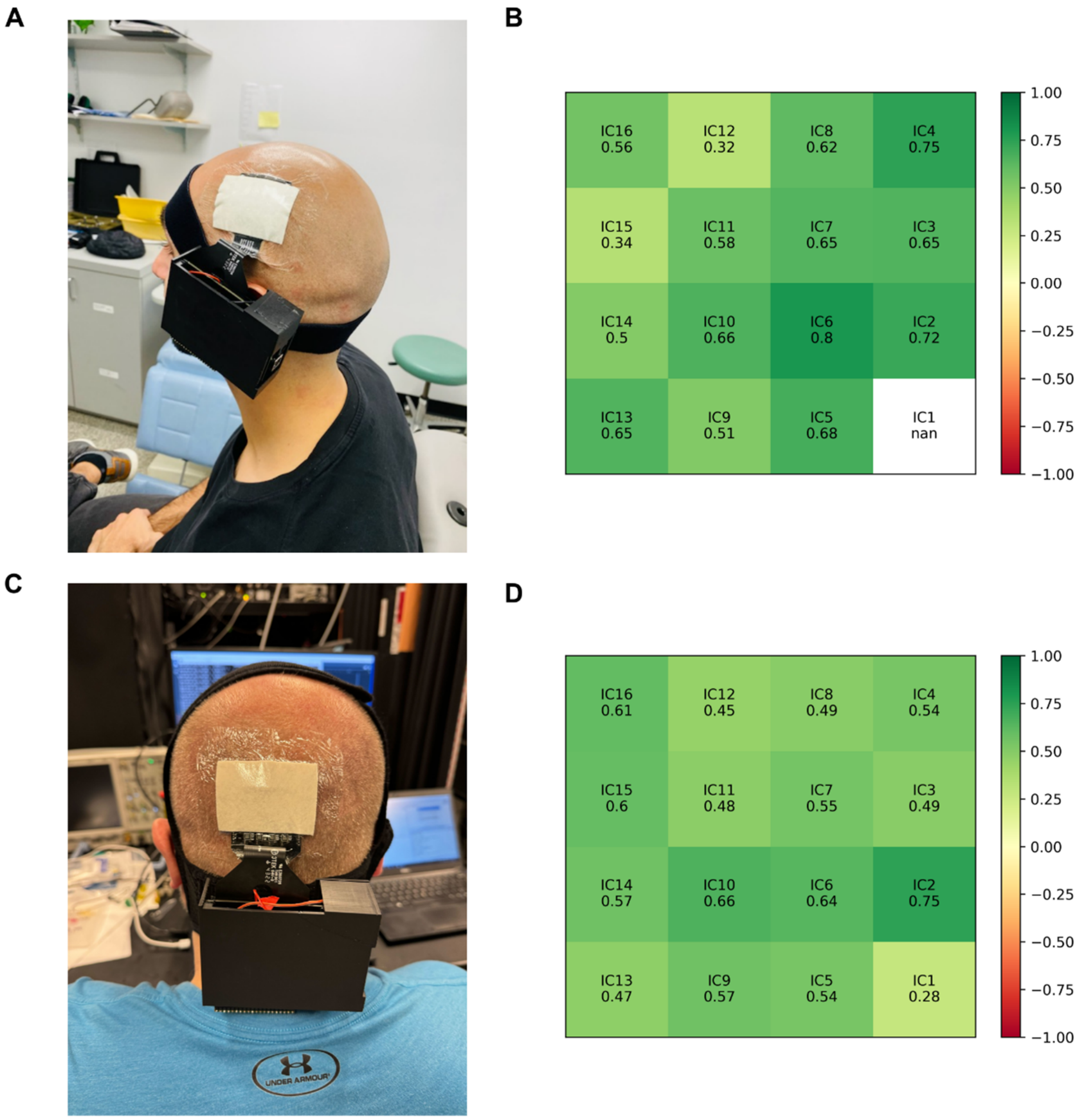


**Fig. S3. Scalp coupling index comparison for different hair types.** (A) Image of the HD-TD-DOT system mounted on the head of a bald subject. Image depicts author Petros D. Petridis. (B) The maximum SCI values for each detector location following device placement over the motor cortex of a bald subject. In this particular case, the device was removed and replaced because of the relatively low SCI values at detectors at IC12 and IC15. IC1's SCI could not be calculated because the 680 nm sources at IC2 and IC5 were not functional. (C) Image of the HD-TD-DOT system mounted on the head of a subject with short, light hair. (D) The maximum SCI values for each detector location following device placement over the visual cortex of a subject with short blond hair. SCI values at each detector location are slightly reduced compared to that of the bald subject, but additional uniformity is provided by a flatter location on the scalp provided by the visual cortex.

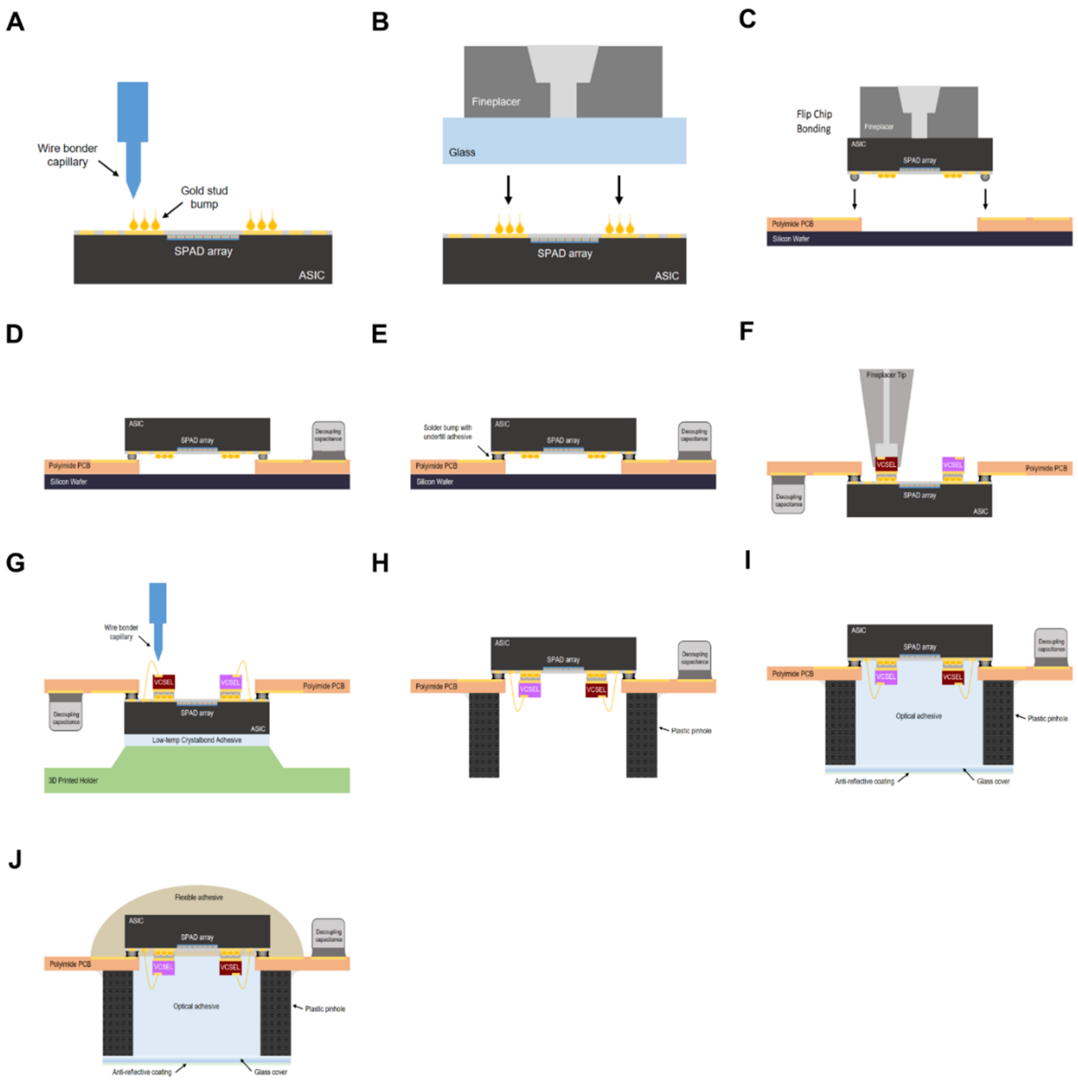


**Fig. S4. Array packaging steps.** (A) Chiplet is stud bumped with gold balls using a wire bonder. (B) Bumps are leveled through application of moderate pressure with a planar piece of glass. (C) ASIC is flip chip bonded to flexible substrate secured down against a silicon wafer for stability. (D) Decoupling capacitance is mounted on flexible substrate and device is placed in reflow oven for solder reflow. (E) Underfill adhesive is dispensed underneath the chiplet to add mechanical stability to solder bump connections. (F) Electrically conductive adhesive is dispensed onto gold stud bumps, and VCSELs are mounted into electrically conductive adhesive before being heat cured to form connection. (G) VCSEL anode pads are wire bonded to the ASIC to form connection with VCSEL driver circuits. A custom 3D printed holder and a small amount of low temperature adhesive allows for securing of the ASIC during the wire bonding process. (H) Plastic pinhole is mounted on the bottom of the flexible substrate. (I) Optical adhesive and glass coverslip with anti-reflective coating are mounted above the chiplet for protection and index matching. This step is optional, and not every flexible device that is prepared will include optical adhesive or a glass coverslip. (J) Flexible, thermally conductive adhesive is dispensed onto the chiplet for protection as well as heatsinking purposes.

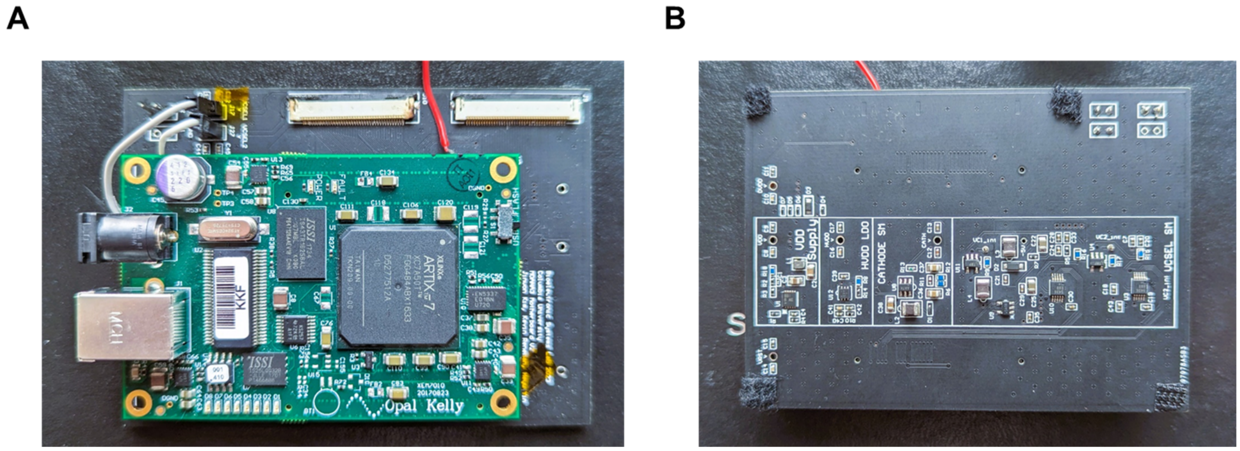


**Fig. S5. Interface Motherboard.** (A) Top of motherboard with FPGA and ZIF connectors. (B) Bottom of motherboard with power conditioning circuitry.

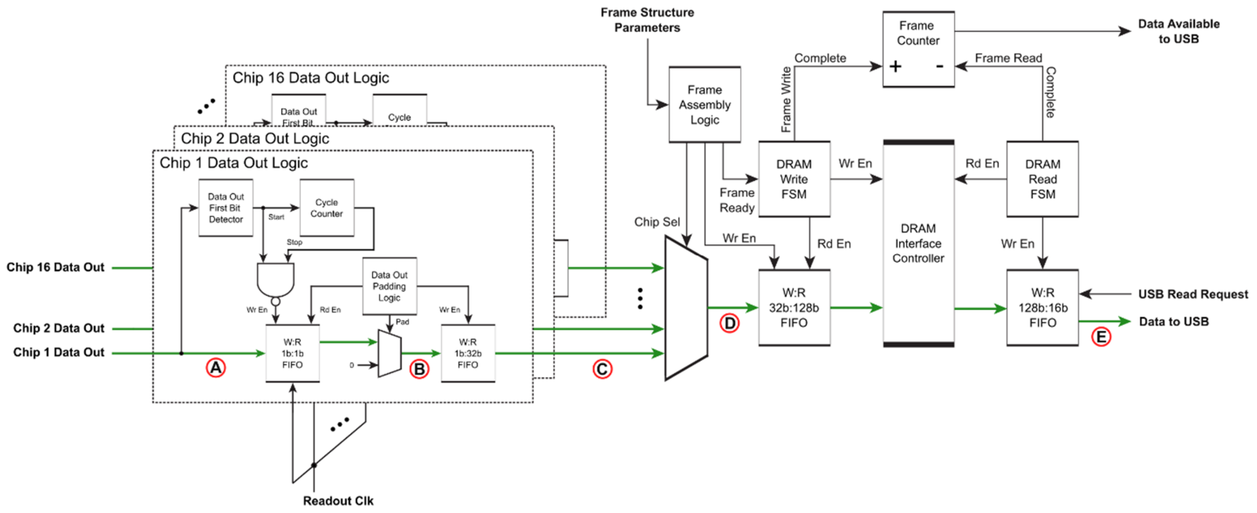


**Fig. S6. FPGA data aggregation block diagram.** The architecture of the logic on the FPGA that handles readout and serialization of data collected from each chiplet on the flexible array. (A) Serialized histogram data flows out from each chip at the completion of a source window. Each chip sends 3000 bits, representing 150 histogram bin values of 20 bits each. (B) Padding logic inserts 12 zeros into the data stream for every 20 bits coming from the chiplet such that histogram data from each ASIC becomes 4-byte aligned. (C) Non-symmetric, first-in first-out (FIFO) memory transforms serial data into 32-bit data. Each 32-bit output from this FIFO corresponds to a single histogram bin value. (D) Frame assembly logic multiplexes data from each chip into a FIFO. Data held in this FIFO represents histogram values from all 16 chiplets collected during a single source window. When all source windows in a frame have been completed, the corresponding data is written into off-chip DRAM. (E) When a configurable number of frames have been written into DRAM (usually at least 10 frames, which is about 3 MB of data), a USB transfer is initiated that moves the specified number of frames to the host PC for visualization and processing.

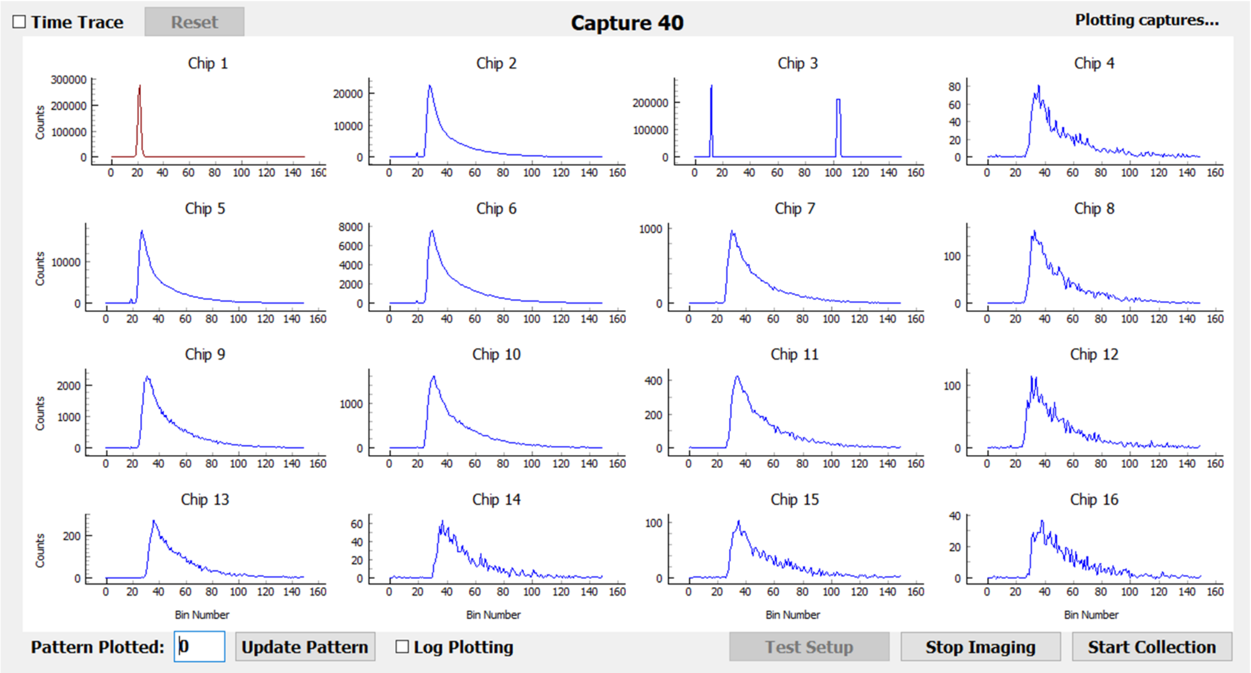


**Fig. S7. Python GUI.** The GUI interface to the system. A test setup is created through the "Test Setup" button and corresponding dialog. The test setup determines such parameters as frame rate, source order within the frame, and integration time per source. ToF histograms can be visualized in real time at each ASIC location in the array. The source location is shown in red, whereas detector locations are shown in blue. By updating the "Pattern Plotted", ToF histogram data for a different source location within the frame can be visualized. Log plotting is also available. The "Time Trace" checkbox plots total photon count history over the course of the experiment at each detector location for visualization of heart rate signals and motion artifacts. The imaging process can be started prior to storage of experimental data so that device warmup can be accommodated.

**A**

| | |
|---|---|
| Detector Responsivity (635 nm) | $2.4 \times 10^{-11}$ $m^2sr$ |
| DNL Variation | <0.12 |
| IRF FWHM | 50%: 505 ps; 10%: 1870 ps; 1%: 7360 ps |
| $R_{AP}$ | 690 nm: 0.3%<br>850 nm: 0.9% |
| Stability (after warm up) | N/No variability < ± 1%<br>FWHM variability < ± 5 ps<br>Mean ToF variability < ± 1 ps |
| Count Rate | 400 MCPS |

**B**

| | VCSEL Driver Setting | VCSEL Cathode Bias (V) | Measured Average Power (μW) | Calcluated Energy per Pulse (pJ) | Measured FWHM (ps) | Calculated Peak Power (mW) |
|---|---|---|---|---|---|---|
| IR VCSEL (850 nm) | 1 | -0.9 | 126.9 | 2.54 | 511 | 4.37 |
| | 2 | -1.0 | 186.2 | 3.72 | 631 | 5.19 |
| | 3 | -1.0 | 226.2 | 4.52 | 706 | 5.64 |
| NIR VCSEL (680 nm) | 3 | -0.4 | 69.4 | 1.39 | 259 | 4.72 |
| | 4 | -0.4 | 136.0 | 2.72 | 381 | 6.28 |
| | 4 | -0.4 | 247.0 | 4.94 | 505 | 8.61 |

**C**

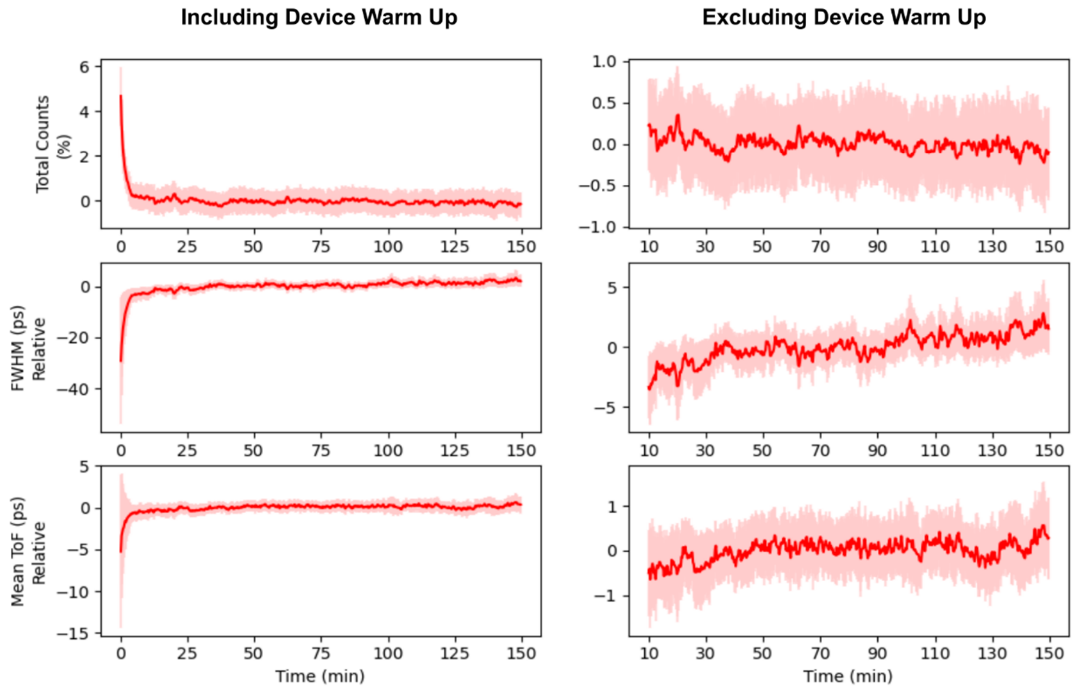


**Fig. S8. Extended BIP results.** (A) BIP metrics and corresponding measured values. (B) Source characterization through measurement of average power and measured FWHM at selected VCSEL driver settings and VCSEL cathode biases. (C) Total counts, FWHM, and mean ToF stability plots for the device over 150 minutes of continuous runtime. Red line represents the mean value, while the surrounding red shading represents plus/minus one standard deviation. Results with and without warm up are presented to make it easier to evaluate the latter portion of the measurement.

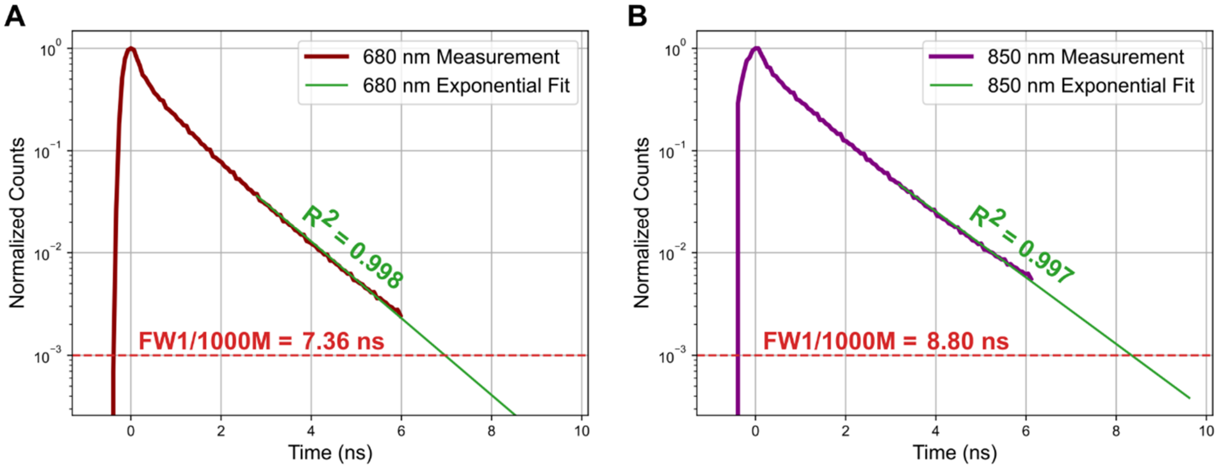


**Fig. S9. Exponential fitting of system IRF to determine FW1/1000M.** (A) Exponential fitting of the tail of the system IRF at 680 nm showing extrapolation of the decaying exponential down to three orders of magnitude below the peak to extract the FW1/1000M. (B) The equivalent plot for the tail of the system IRF at 850 nm.

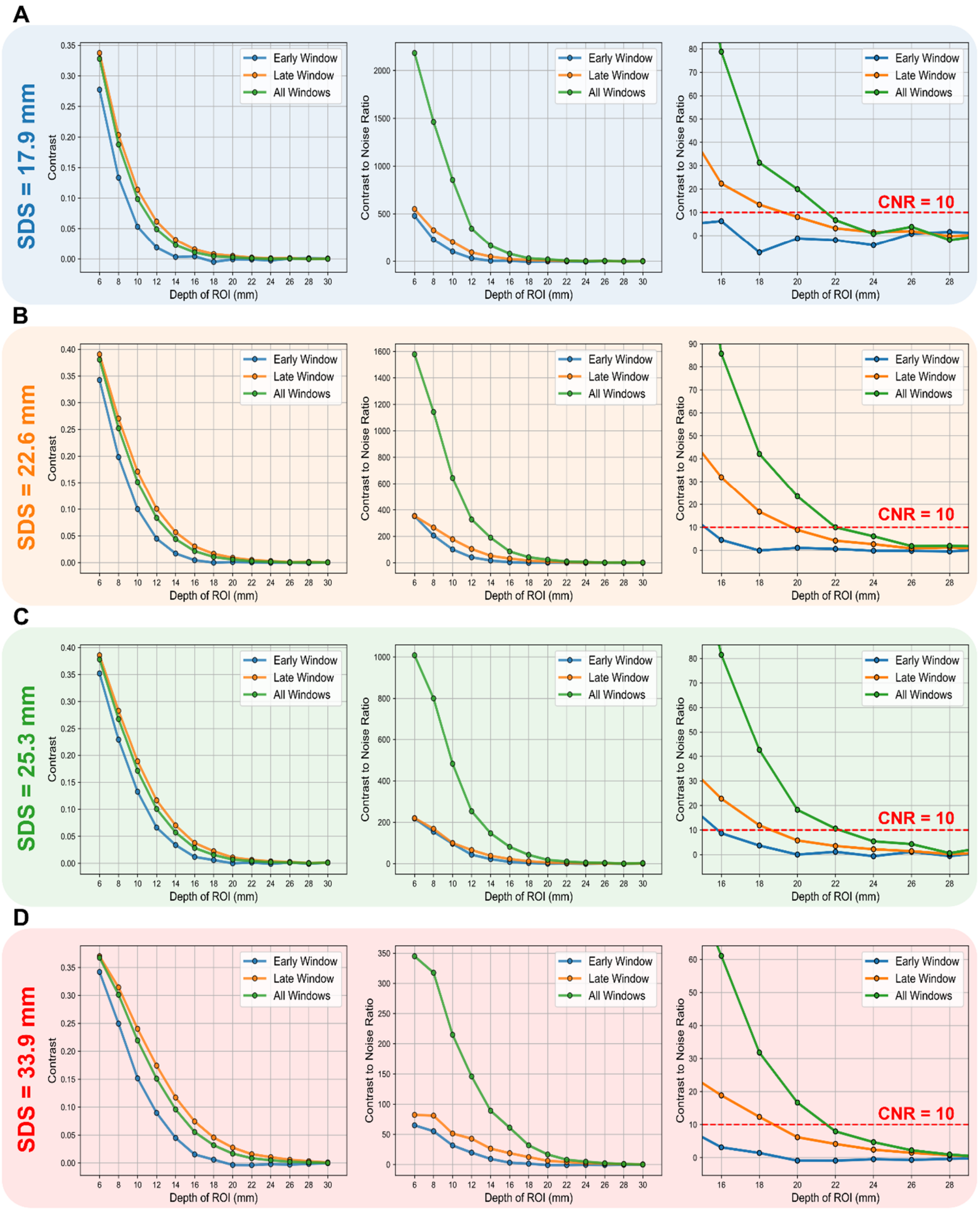


**Fig. S10. Extended contrast and contrast-to-noise-ratio results.** Contrast (left column), contrast-to-noise ratio (center column), and contrast-to-noise ratio for depths beyond 16 mm (right column) for source-detector separations of (A) 17.9 mm, (B) 22.6 mm, (C) 25.3 mm, and (D) 33.9 mm. Contrast and contrast-to-noise results for 28.9 mm source-detector separation are presented in **Fig 3E** and **Fig 3F**, respectively.

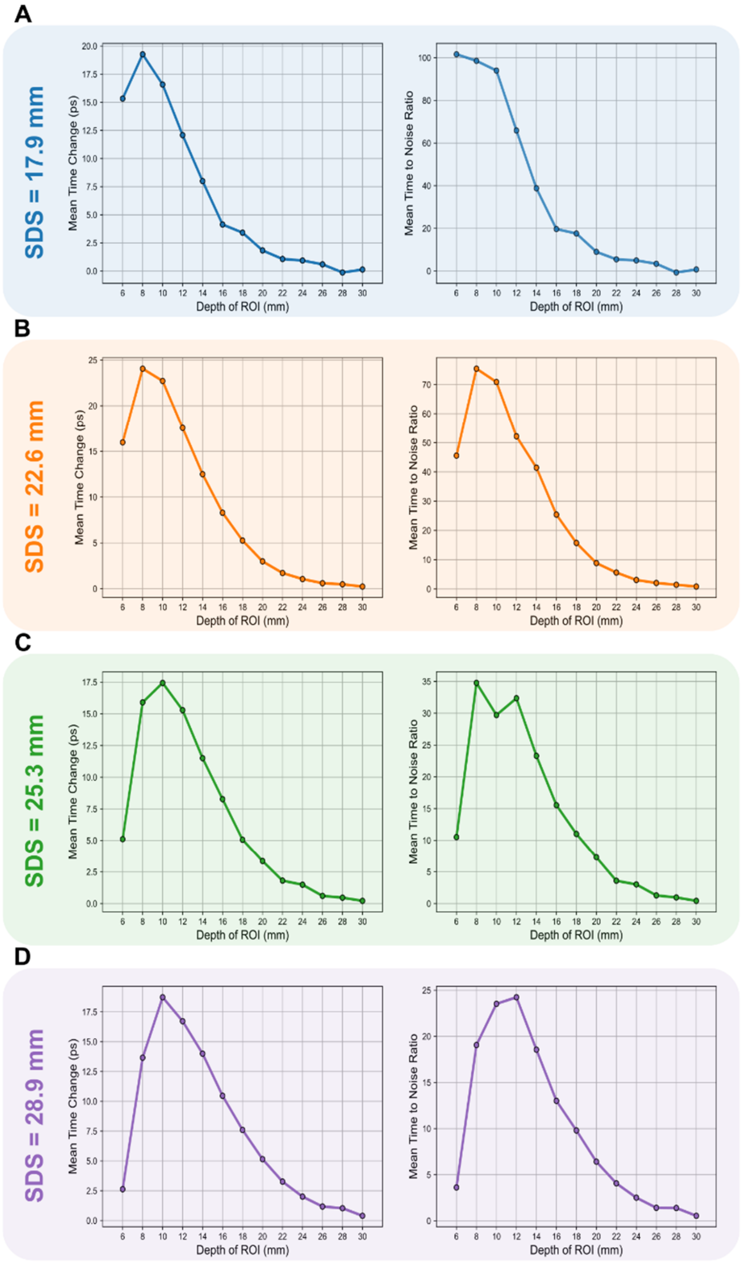


**Fig. S11. Extended mean-time results.** Mean time change (left column) and mean-time-to-noise ratio (right column) at source-detector separations of (A) 17.9 mm, (B) 22.6 mm, (C) 25.3 mm, and (D) 28.9 mm. Mean time change and mean-time-to-noise results for 33.9 mm source-detector separation are not presented because the mean-time-to-noise ratio was too low.

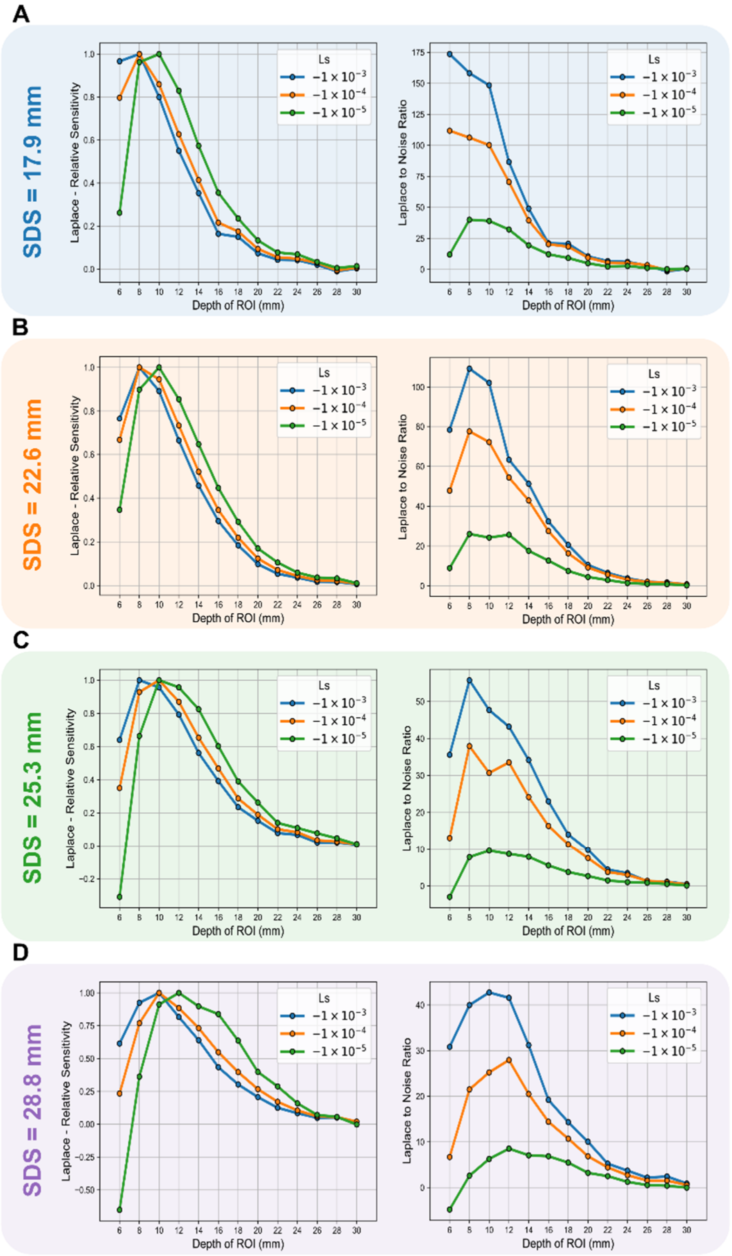


**Fig. S12. Extended Laplace results.** Laplace normalized sensitivity (left column) and Laplace-to-noise ratio (right column) for three different selections of exponential coefficient at source-detector separations of (A) 17.9 mm, (B) 22.6 mm, (C) 25.3 mm, and (D) 28.9 mm. Laplace normalized sensitivity and Laplace-to-noise results for 33.9 mm source-detector separation are not presented because the Laplace-to-noise ratio was too low.

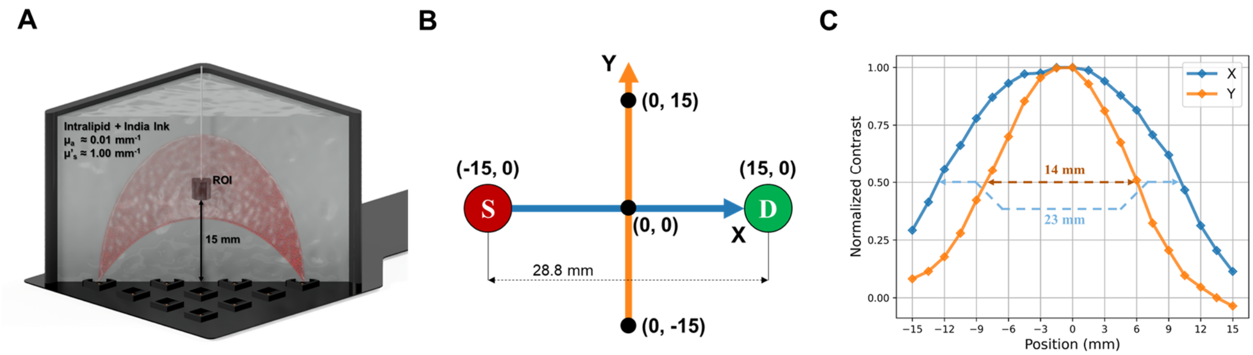


**Fig. S13. nEUROPt lateral spatial resolution results.** (A) Experiment setup showing an ROI suspended in a tank filled with Intralipid and India ink at a depth of15 mm. (B) Illustration of the ROI movement along the X and Y axis and how X and Y positions are defined relative to the source and detector position. (C) Experiment results showing lateral resolution of 23 mm along the X axis and 14 mm along the Y axis.

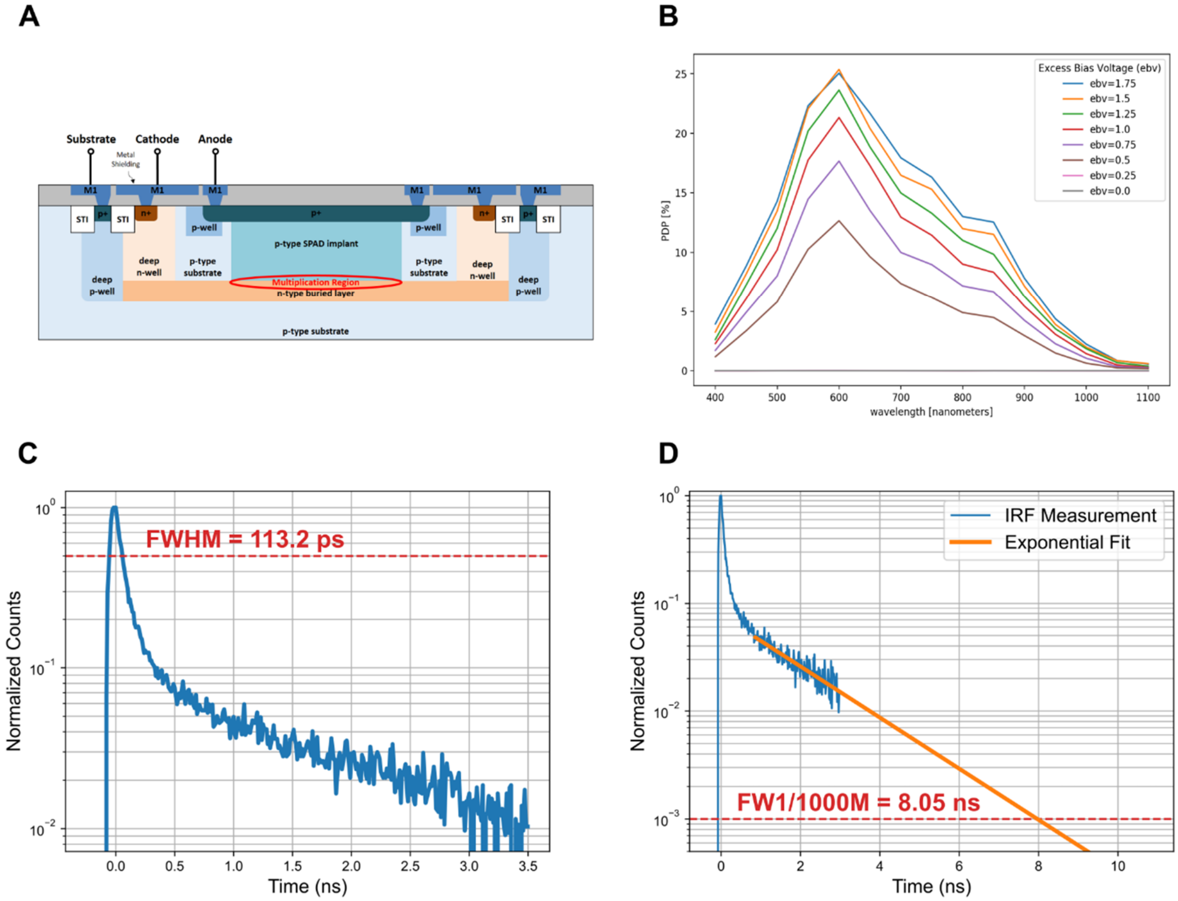


**Fig. S14. SPAD structure and characterization.** (A) SPAD cross-section with electrical terminals shown and photosensitive multiplication region indicated by red ellipse. (B) SPAD PDP at different wavelengths for various excess bias voltages (EBV). (C) SPAD IRF measurement showing the FWHM at 800 nm. (D) SPAD IRF measurement and exponential fit line used to extract the FW1/1000M of the SPAD at 800 nm. $R^2$ for the exponential fit is 0.77.

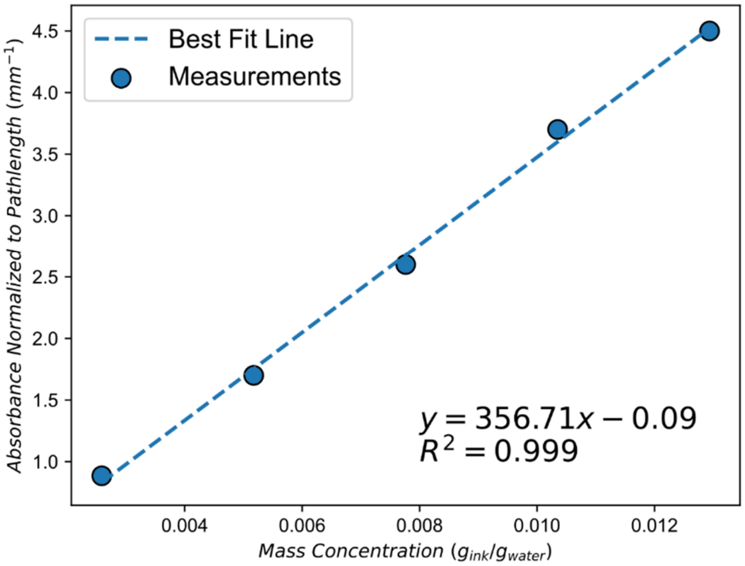


**Fig. S15. Optical absorption characteristics of India ink.** India ink is used in the creation of optical phantoms in this work. It has been shown that the absorption coefficient of India ink varies significantly from batch to batch. As a result, it is necessary to characterize the India ink before it can be correctly used to realize an absorption coefficient in an optical phantom. A Thermofisher NanoDrop 1000 Spectrophotometer was used to obtain this measurement data.

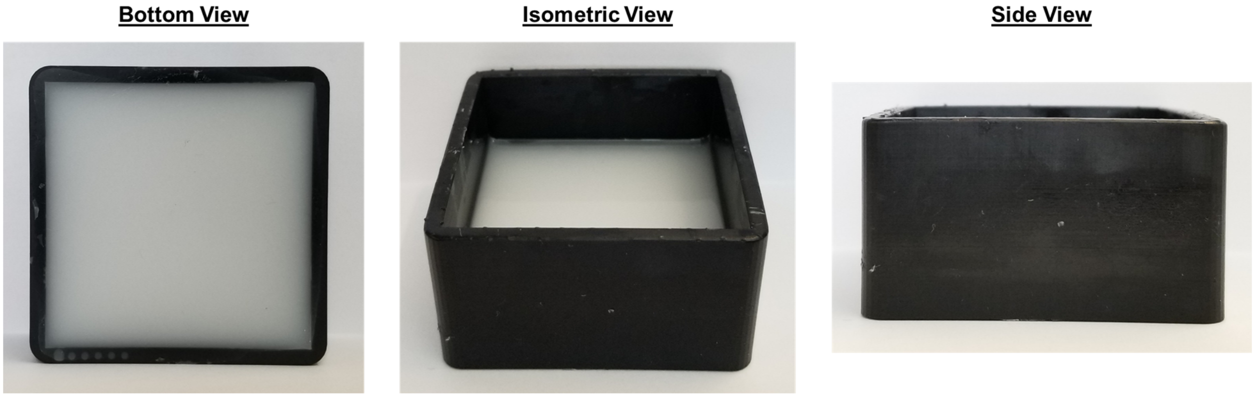


**Fig. S16. Visual cortex phantom.** Bottom, isometric, and side view of the visual cortex phantom.

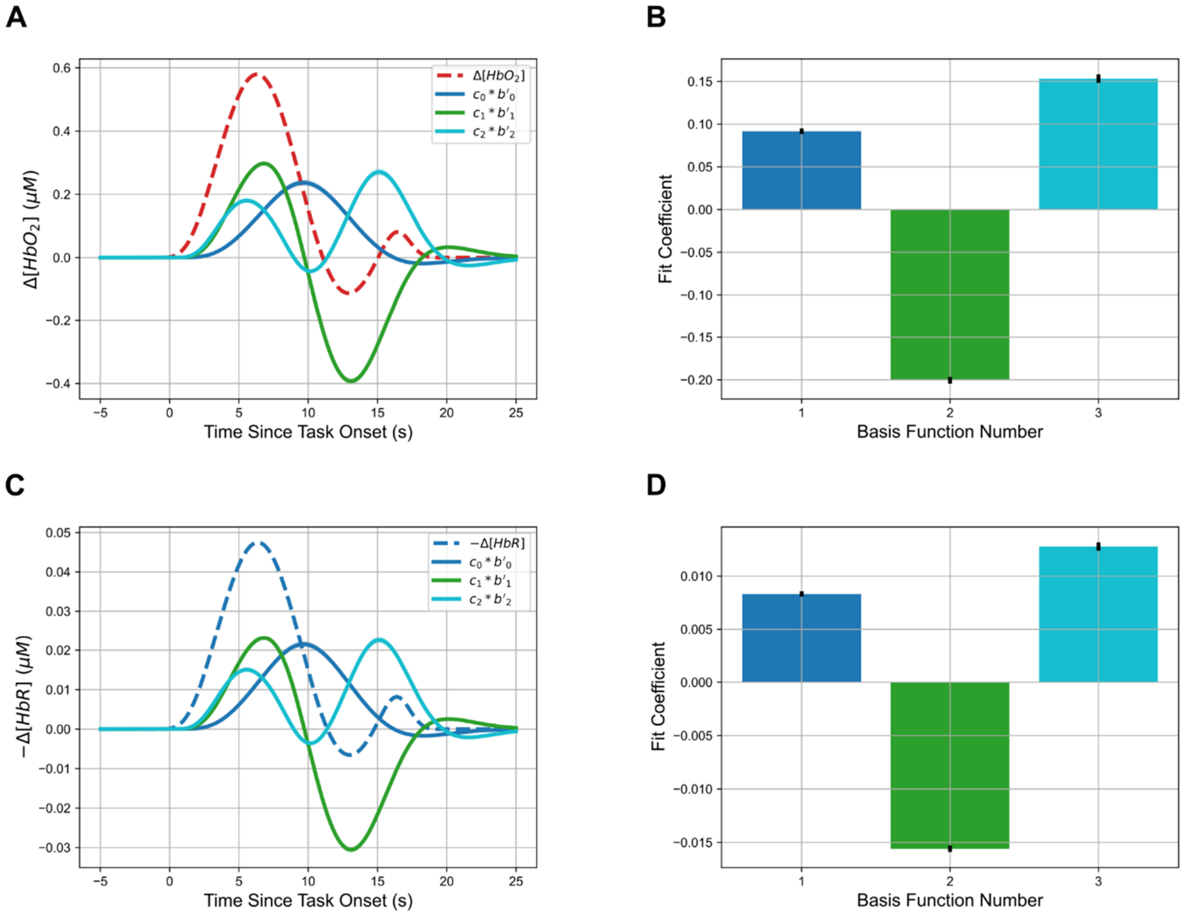


**Fig. S17. Basis function fits.** (A) [$HbO_2$] time trace and resulting basis function fit. Each basis function ($b_n'$), which has been convolved with a boxcar that has value 1 during the 6 s period corresponding to the finger tapping task, is shown multiplied by its corresponding fit coefficient ($c_n$). (B) [$HbO_2$] fit coefficients by basis function number. (C) Inverted [HbR] time trace and resulting basis function fit. Each basis function ($b_n'$), which has been convolved with a boxcar that has value 1 during the 6 s period corresponding to the finger tapping task, is shown multiplied by its corresponding fit coefficient ($c_n$). (D) [HbR] fit coefficients by basis function number.

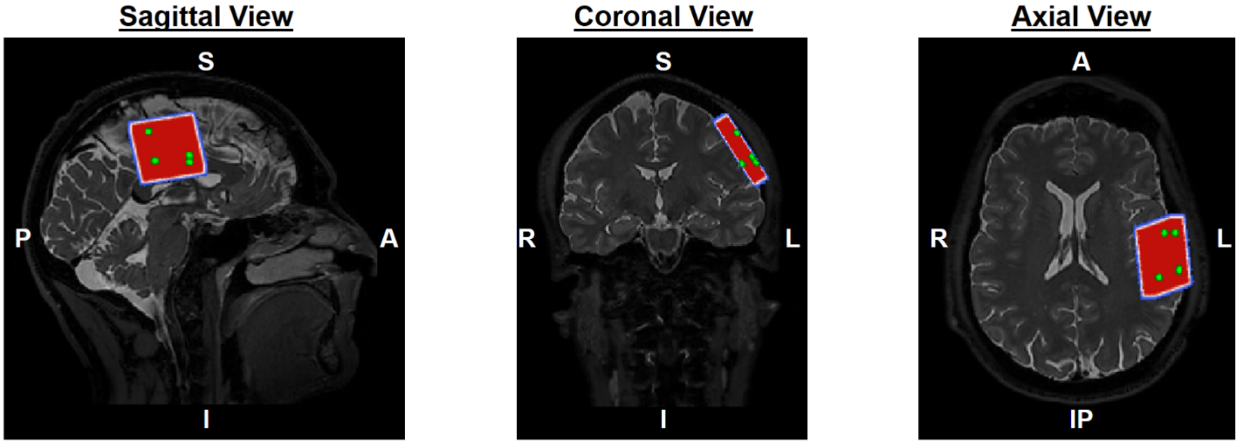


**Fig. S18. Alignment of reconstruction volume with T2 in FSLEyes.** An illustration of the HD-TD-DOT array after registration with the T2 volume. The reconstruction volume is shown in red, and is displayed as a max intensity projection to give an impression of the alignment in 3D space. The green voxels within the reconstruction volume are the alignment points, 3 of them corresponding to the corners of the array at z = 16 mm, and 1 of them corresponding to the corner of the array at z = 30 mm.

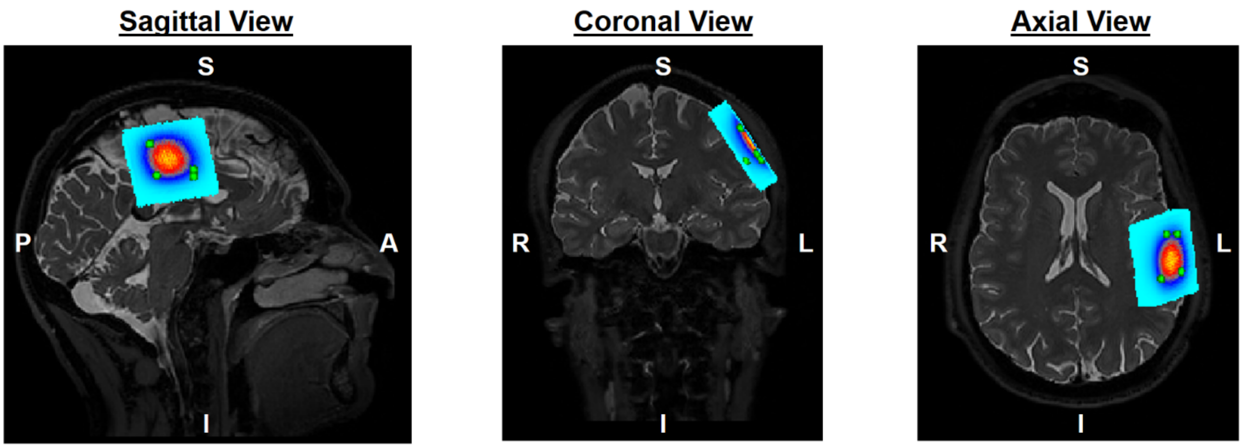


**Fig. S19. $HbO_2$ reconstruction volume after alignment with T2 in FSLEyes.** An illustration of the $HbO_2$ changes after alignment with the T2. The Render3 colormap is used here. The green voxels within the reconstruction volume are the alignment points.

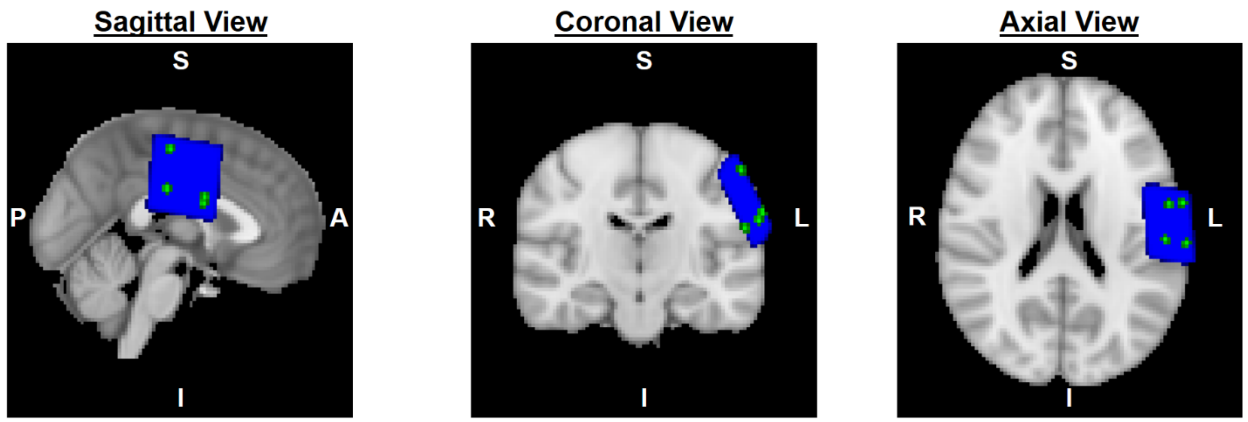


**Fig. S20. T2 and reconstruction volumes in Montreal Neurological Institute (MNI) space.** An illustration of the reconstruction volume and brain extracted from T2 scan following MNI registration. The reconstruction volume is shown in blue, and is displayed as a max intensity projection. The green voxels within the reconstruction volume are the alignment points.

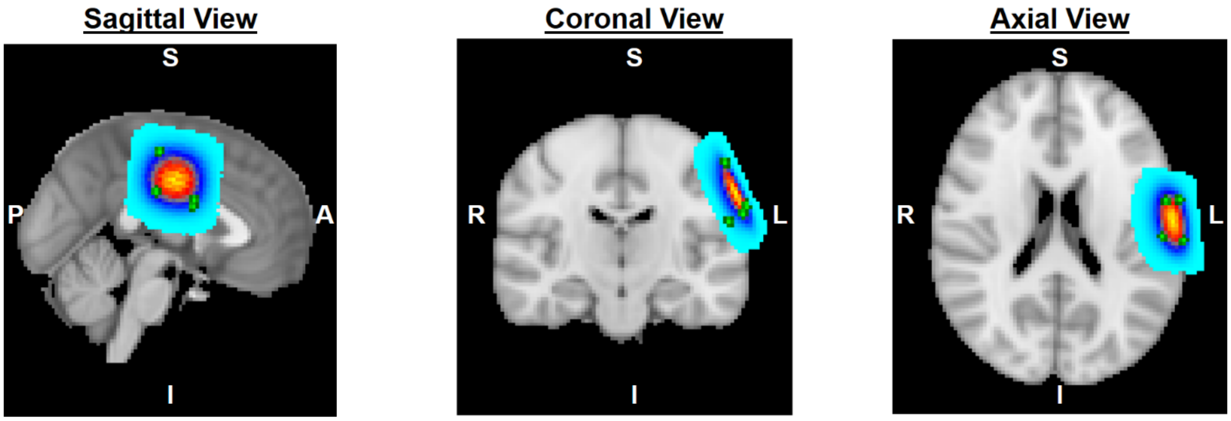


**Fig. S21. T2 and $HbO_2$ reconstruction volumes in MNI Space.** An illustration of the [$HbO_2$] changes after MNI registration. The Render3 colormap is used here. The green voxels within the reconstruction volume are the alignment points.

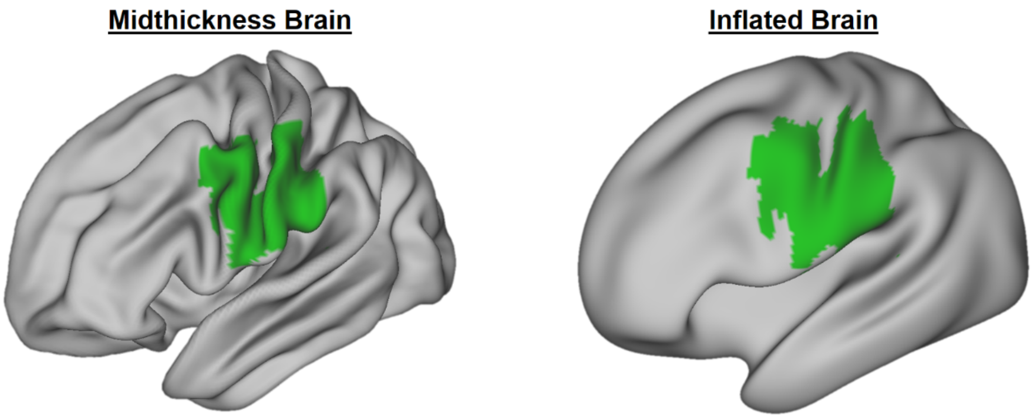


**Fig. S22. Device field-of-view following volume-to-surface mapping.** The field-of-view of the HD-TD-DOT array is shown in green on the midthickness brain (left) and on the inflated brain (right).

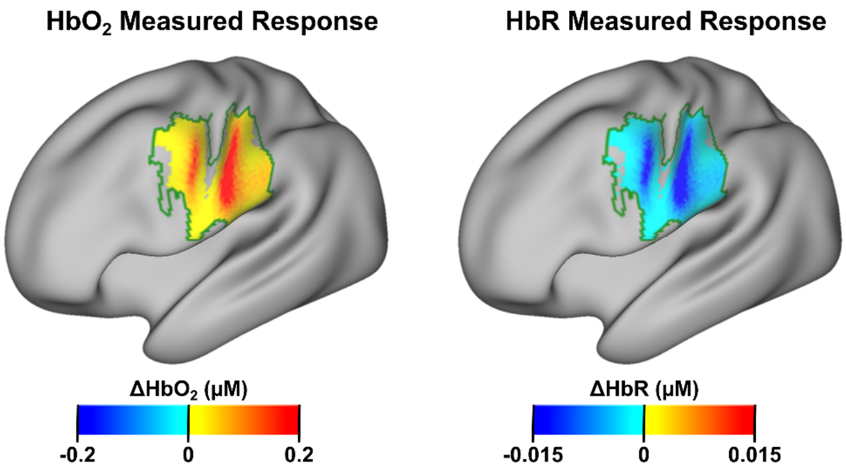


**Fig. S23. Chromophore Concentration Maps Following Volume to Surface Mapping.** **Δ[**$HbO_2$] (left) and Δ[HbR] (right) measured by the Micro-DOT device during the finger tapping task.

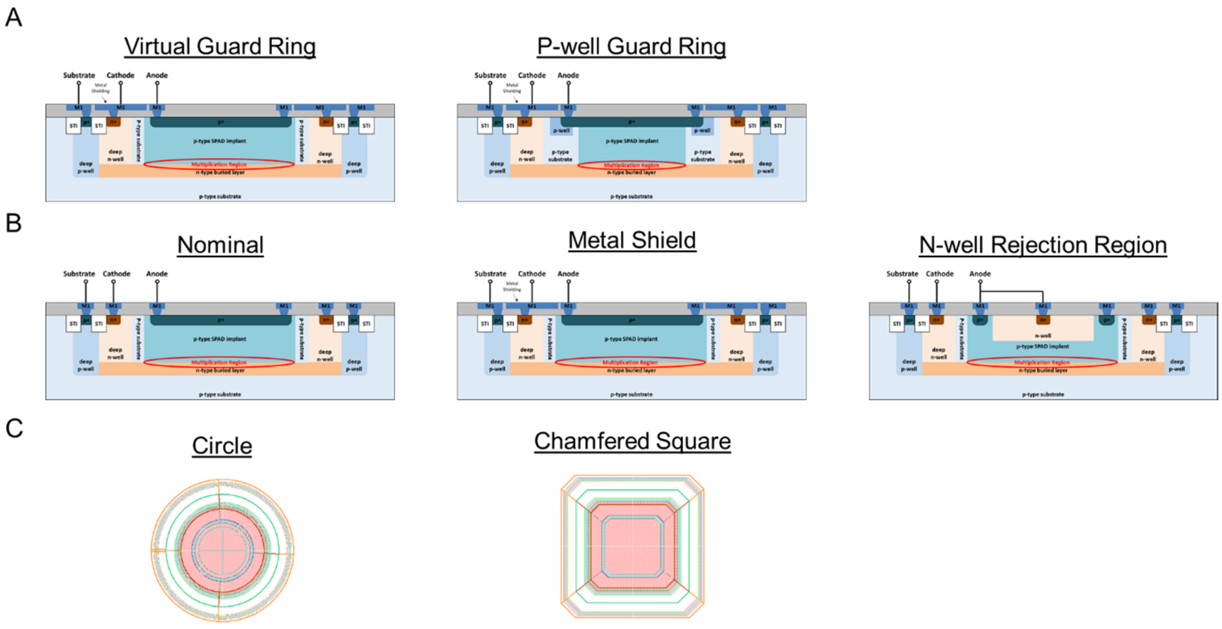


**Fig. S24. SPAD variations in the DOE array.** (A) Cross-section view of layer structure variations showing two different guard rings structures. (B) Cross-section view of metal shield and n-well rejection region variations. (C) Shape variations. These images show the footprint of the SPAD looking perpendicular to the surface of the silicon.

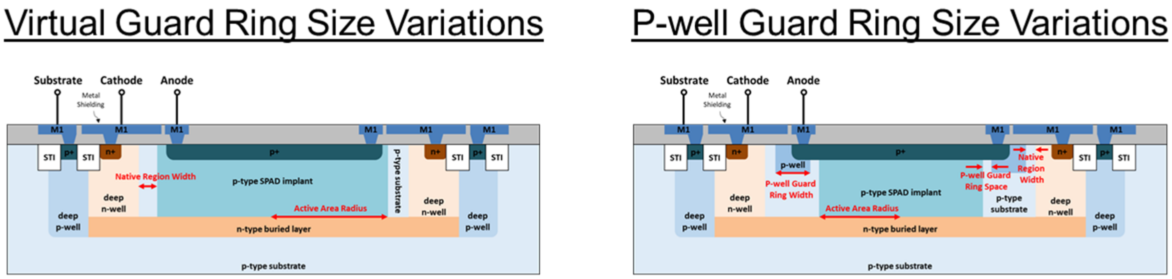


**Fig. S25. Size variations in the DOE array.** Size variations for each guard-ring type.

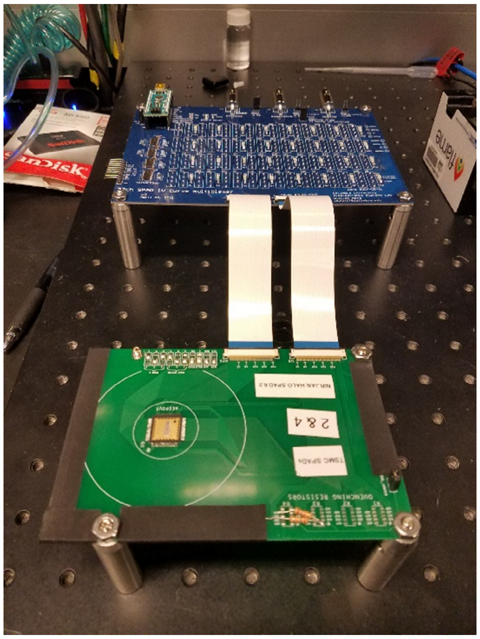

**Fig. S26. IV-curve measurement setup.** Image of automated IV curve measurement system showing the motherboard (top of image) and daughterboard (bottom of image).

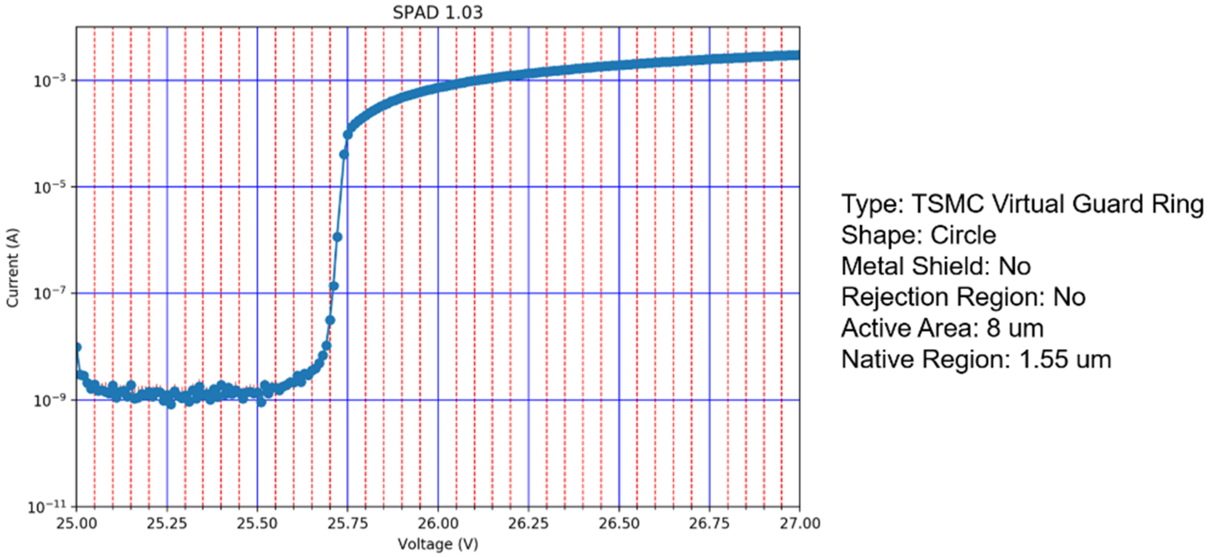


**Fig. S27. Automated IV curve output.** Output figure generated by the automated IV curve setup following IV sweep. The plot output also includes relevant structural details such as guard ring type, shape, and active area diameter.

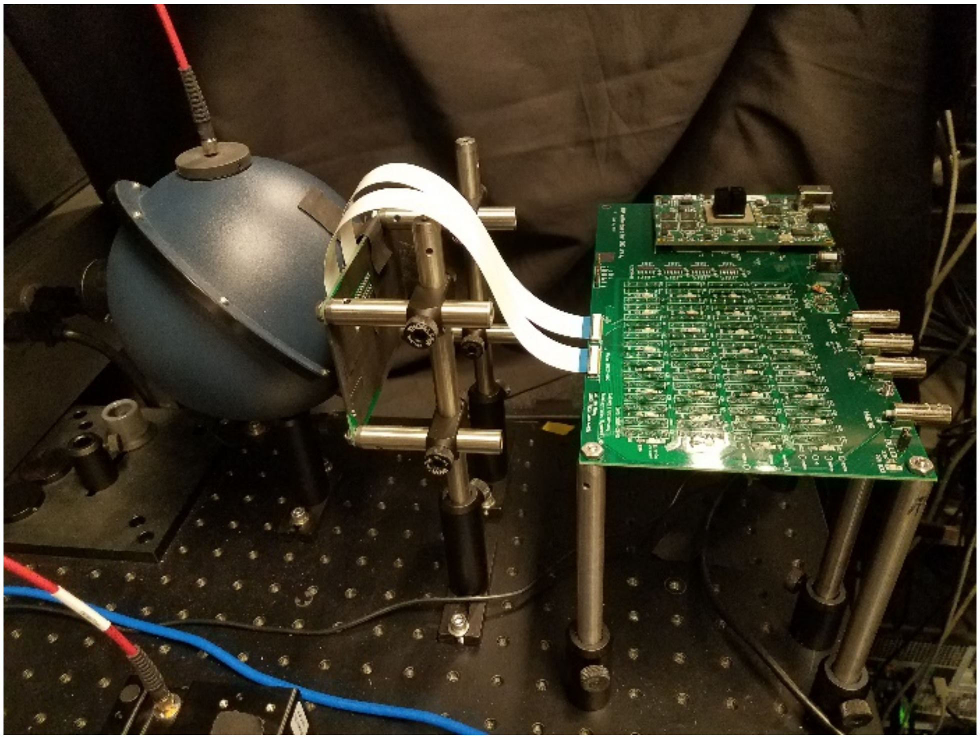

**Fig. S28. Automated PDP/DCR testing setup.** Motherboard (right side) connected to daughterboard mounted into port of integrating sphere for PDP/DCR measurement.

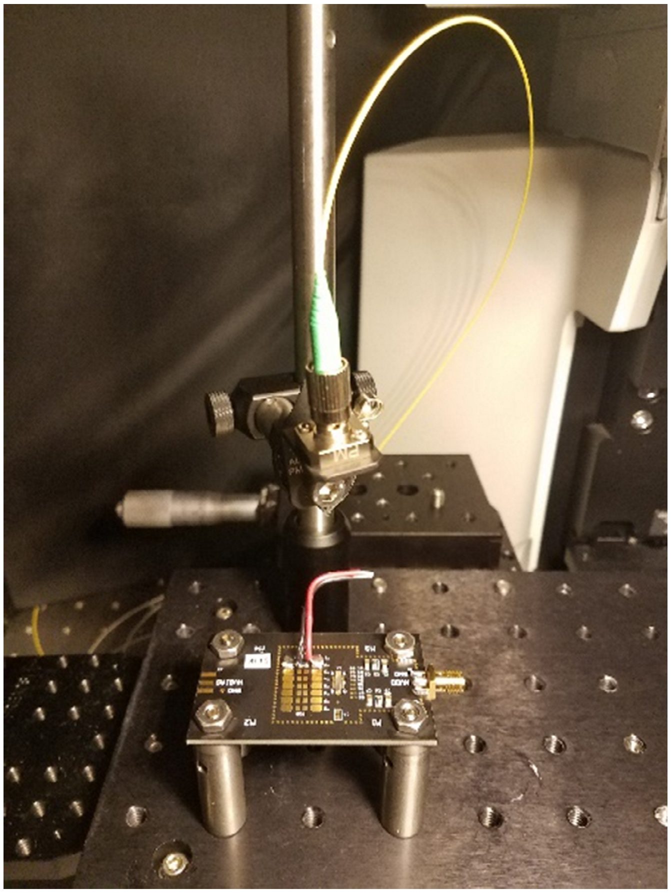

**Fig. S29. IRF testing setup.** Overhead mounted fiber for coupling pulsed laser to SPAD. Connection to oscilloscope not shown.

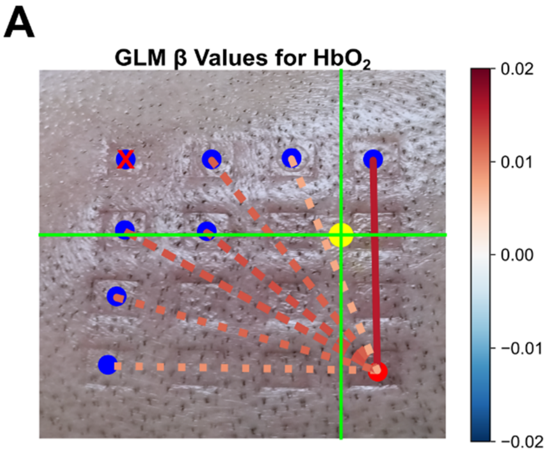


**Fig. S30. 6-s finger tapping GLM statistics.** (A) GLM results for [$HbO_2$]. Solid lines indicate statistical significant ($p < 0.05$) while dashed lines represent statistical non-significance ($p \geq 0.05$). (B) GLM results for [HbR].

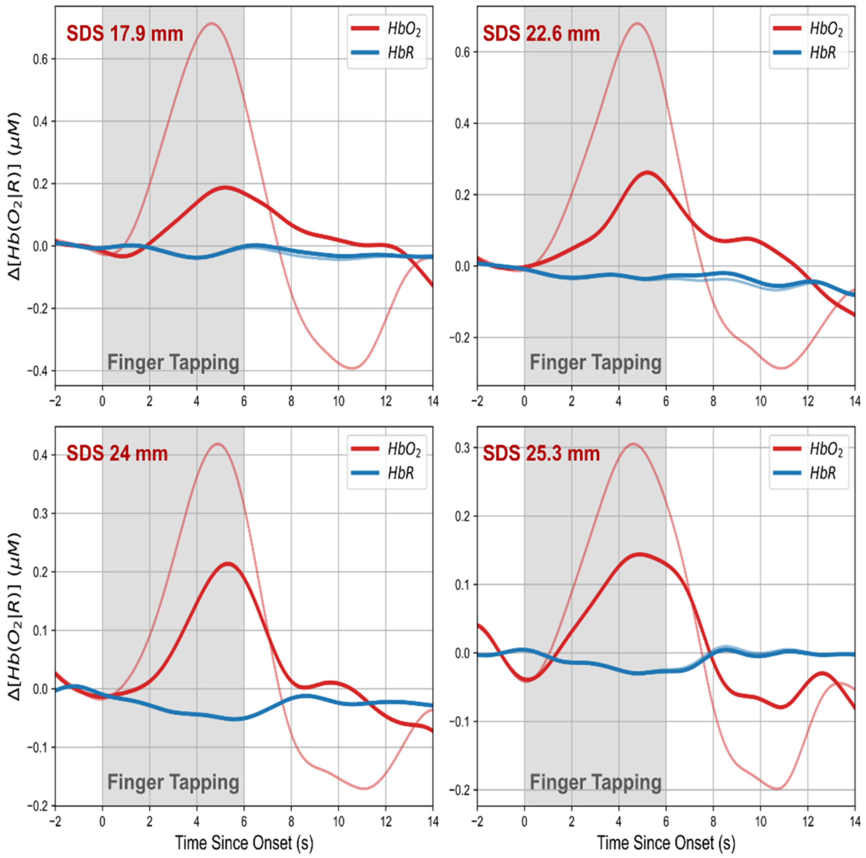


**Fig. S31. 6-s finger tapping block averages after regression of short channel data.** Block averaged time traces of the *E* feature for various SDSs after regressing out shallow activation captured by a channel with 8-mm SDS are shown. Red traces represent [$HbO_2$] while blue traces represent [HbR]. The block average time traces before regressing out shallow activation are shown in a lighter color for comparison.

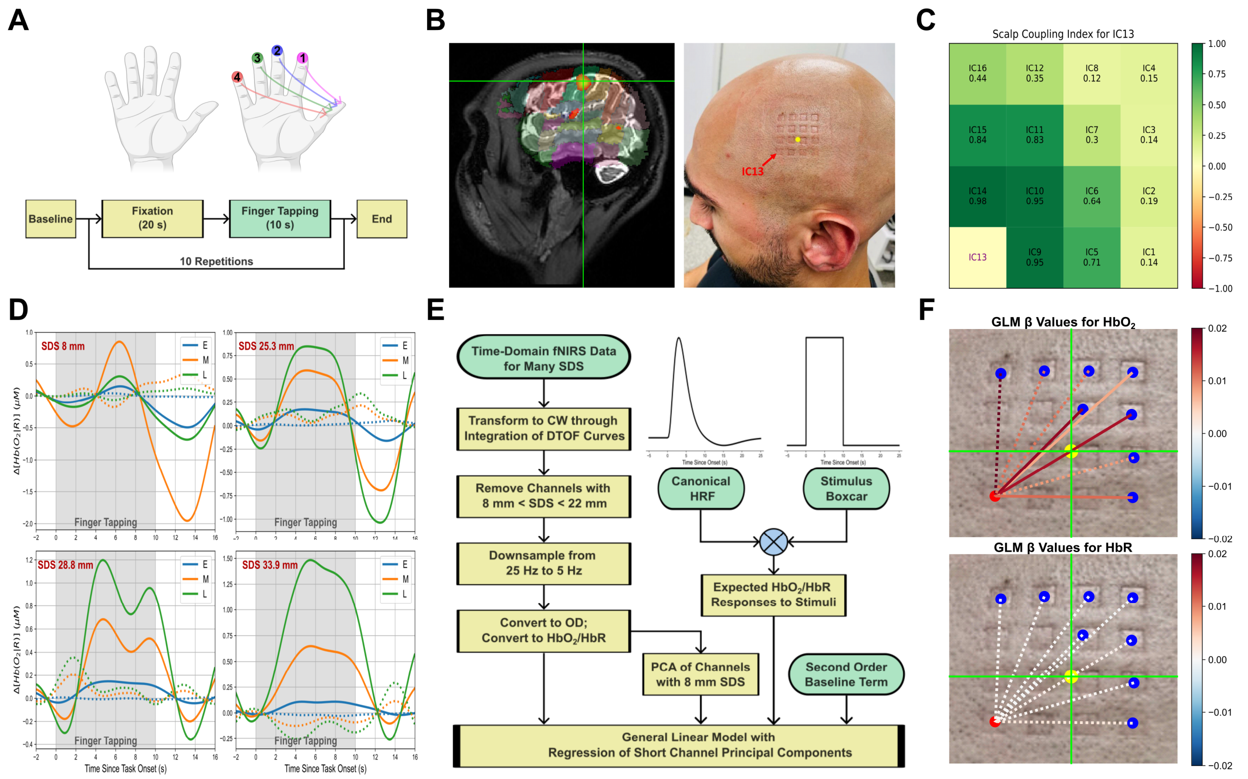


**Fig. S32. 10-s finger-tapping results.** (A) Finger tapping protocol with 10-s duration. (B) fMRI localized activation during finger tapping task and corresponding array placement. The yellow dot in the image on the right represents the activated region localized with fMRI. Image depicts author Petros D. Petridis. (C) Scalp coupling index for a single source. (D) Block averages for four different channels of varied SDS in response to the task. (E) GLM data processing block diagram. (F) GLM results for [HbO$_2$] (upper) and [HbR] (lower). Solid lines indicate statistical significant ($p < 0.05$) while dashed lines represent statistical non-significance ($p \geq 0.05$).

| | | *Milan Probe SiPM*[19] | *MAESTROS*[20] | *Kernel Flow*[21] | ***Micro-DOT (this work)*** |
|---|---|---|---|---|---|
| ***System*** | **Technology** | Detection Probe with External Lasers | Equipment Cart | Rigid Module | **Flexible Patch** |
| | **Source Type** | Laser Diode Module | Supercontinuum Laser | Edge-Emitting Laser | **VCSEL** |
| | **Wavelengths (nm)** | 670 & 830 | 650-900 (adjustable) | 690 & 850 | **680 & 850** |
| | **Number of Source Locations** | 1 | 1 | 52 | **16** |
| | **Detector Type** | Silicon Photomultiplier | Photomultiplier Tubes | SPAD Array | **SPAD Array** |
| | **Number of Detectors** | 1 | 4 | 312 | **16** |
| | **Number of Channels** | 1 | 4 | 2206 | **256** |
| | **Shortest Channel (mm)** | 28.8 | N/R | 10 | **0.4** |
| | **Longest Channel (mm)** | 30.1 | N/R | 60 | **33.9** |
| | **Field-of-View** | N/A | N/A | Entire Head | **4 cm × 4 cm** |
| | **Total Mass (g)** | N/R | N/A | 2050 | **96** |
| | **Total Power (W)** | N/R | N/R | $50^{+}$ | **3.4** |
| ***BIP*** | **Stability (after warm up)** | N/No < ± 0.5%<br>FWHM < ± 10 ps<br>Mean ToF ± 2 ps | N/No < ± 1%<br>FWHM N/R<br>Mean ToF ± 5 ps | N/No < ± 2%<br>FWHM < ± 20 ps<br>Mean ToF ± 2 ps | **N/No < ± 1%<br>FWHM < ± 5 ps<br>Mean ToF < ± 1 ps** |
| | **Afterpulsing Probability ($R_{AP}$)** | N/R | 0.15% | 690 nm: 0.7%<br>850 nm: 0.5% | **690 nm: 0.3%<br>850 nm: 0.9%** |
| | **Detector Responsivity ($m^2sr$)** | $3.04\times10^{-8}$ - $3.3\times10^{-8}$ | $2\times10^{-8}$ | $7.2\times10^{-9}$ | **$2.4\times10^{-11}$** |
| | **System IRF FWHM (ps)** | 308-556 | 465 | 290-350 | **680 nm: 505<br>850 nm: 706** |
| | **System IRF FW1/1000M (ps)** | N/R | 3250 | 680 nm: 1560<br>850 nm: 1620 | **680 nm: 7360<br>850 nm: 8800** |
| | **Peak Source Power (mW)** | $145^{\dagger}$ | N/R (3.4 mW average) | $37^{\dagger}$ | **680 nm: 8.61<br>850 nm: 5.64** |
| | **DNL Variation** | 0.03 | N/R | < 0.5 | **<0.12** |
| | **Count Rate (MCPS)** | 40 | N/R | >1500 | **400** |
| ***nEUROPt*** | **Contrast at 1.6 cm for Integration Time of 1-s** | 0.4 | 0.125 | 0.2 | **0.02** |
| | **CNR at 1.6 cm Depth for Integration Time of 1-s** | ~500 | N/R | $200^{*}$ | **65** |
| | **MNR at 1.6 cm Depth for Integration Time of 1-s** | N/R | N/R | $20^{*}$ | **13** |

$^{\dagger}$ Estimated; $^{+}$ Max, includes IMU and EEG sensors; $^{*}$ From Kernel Flow 2; N/R: Not Reported; N/A Not Applicable.

**Table S1. Time-domain fNIRS system comparison.** System specifications, BIP results, and nEUROPt results for state-of-the-art time-domain fNIRS systems for comparison against the micro-DOT system.

| Region | $\mu_a$ ($mm^{-1}$) | $\mu_s'$ ($mm^{-1}$) |
|---|---|---|
| Scalp | 0.02 | 0.70 |
| Skull | 0.02 | 0.90 |
| CSF | 0.003 | 0.125 |
| GM | 0.02 | 0.75 |

**Table S2. Visual-cortex-phantom optical properties.** The optical properties of each of the four layers of the visual cortex phantom are shown. The scalp, skull, and CSF are all solid PDMS layers created by mixing titanium dioxide ($TiO_2$) particles and India ink into the PDMS before curing. The grey matter (GM) layer is a liquid layer created by creating a dilution of Intralipid 20% emulsion and India ink in distilled water.

| SPAD | 1.01 | 1.03 | 1.05 | 2.03 | 2.05 | 3.01 | 4.03 | 5.01 | 8.01 |
|---|---|---|---|---|---|---|---|---|---|
| **Guard Ring Type** | Virtual | Virtual | Virtual | Virtual | Virtual | Virtual | Virtual | Virtual | P-Well |
| **SPAD Shape** | Circle | Circle | Circle | Circle | Circle | Circle | Square | Square | Circle |
| **AA Radius (um)** | 4.5 | 5.5 | 4.5 | 6.5 | 4.5 | 4.5 | 6.5 | 4.5 | 5.45 |
| **NR Width (um)** | 3.05 | 4.05 | 4.05 | 3.05 | 4.05 | 3.05 | 3.05 | 3.05 | 3.05 |
| **PWG Space (um)** | - | - | - | - | - | - | - | - | 0.25 |
| **PWG Width (um)** | - | - | - | - | - | - | - | - | 0.7 |
| **N-Well Reject** | No | No | No | No | No | Yes | No | No | No |
| **Metal Shield** | No | No | No | Yes | Yes | Yes | No | Yes | No |
| **PDP @650nm (%)** | 18.95 | 27.44 | 19.06 | 18.92 | 17.62 | 8.1 | 20.95 | 17.68 | 13.15 |
| **PDP @850nm (%)** | 9.87 | 14.33 | 10.64 | 9.2 | 8.1 | 5.8 | 12.21 | 8.5 | 5.59 |
| **DCR (Hz)** | 18.2 | 31.4 | 14.7 | 40 | 18.6 | 37 | 36.3 | 27.5 | 1366.1 |
| **Jitter (ps)** | 182.1 | 223.9 | 159.9 | 125.9 | 115.5 | 82.5 | 131.9 | 115.5 | 102 |

| SPAD | 9.01 | 9.05 | 9.08 | 11.01 | 11.02 | 11.03 | 11.05 | 12.01 |
|---|---|---|---|---|---|---|---|---|
| **Guard Ring Type** | P-Well | P-Well | P-Well | P-Well | P-Well | P-Well | P-Well | P-Well |
| **SPAD Shape** | Circle | Circle | Circle | Square | Square | Square | Square | Square |
| **AA Radius (um)** | 5.45 | 5.45 | 5.95 | 5.45 | 3.45 | 7.45 | 5.45 | 5.45 |
| **NR Width (um)** | 3.05 | 4.05 | 3.05 | 3.05 | 3.05 | 3.05 | 4.05 | 3.05 |
| **PWG Space (um)** | 0.25 | 0.25 | 0.75 | 0.25 | 0.25 | 0.25 | 0.25 | 0.25 |
| **PWG Width (um)** | 0.7 | 0.7 | 0.7 | 0.7 | 0.7 | 0.7 | 0.7 | 0.7 |
| **N-Well Reject** | No | No | No | No | No | No | No | No |
| **Metal Shield** | Yes | Yes | Yes | No | No | No | No | Yes |
| **PDP @650nm (%)** | 19.45 | 21.71 | 18.25 | 18.21 | 20.14 | 20.42 | 15.79 | 20.18 |
| **PDP @850nm (%)** | 9.55 | 10.51 | 9.32 | 9.56 | 9.87 | 11.5 | 7.99 | 9.05 |
| **DCR (Hz)** | 32.4 | 6.9 | 23.1 | 289.3 | 6.6 | 28.3 | 18 | 46.1 |
| **Jitter (ps)** | 143.9 | 115.5 | 108.8 | 183.9 | 120 | 92 | 125.7 | 145.9 |

**Table S3. DOE-array characterization results.** Complete characterization results for the SPAD structures in the DOE array. Relevant structural parameters as well as performance metrics are included.
